# Identifying Functional Homologues in Human and Marmoset Brain Networks via Movie-Driven Ultra-High Field fMRI

**DOI:** 10.1101/2024.09.05.611482

**Authors:** A. Zanini, A. Dureux, R.S. Menon, S. Everling

**Affiliations:** Centre for Functional and Metabolic Mapping, Robarts Research Institute, University of Western Ontario, London, Ontario, Canada; Department of Physiology and Pharmacology, University of Western Ontario, London, Ontario, Canada

**Keywords:** Functional MRI, marmoset, comparative neuroimaging, cortical mapping

## Abstract

Numerous task-based functional magnetic resonance imaging (fMRI) studies have demonstrated that complex neural functions such as language processing, action observation, face recognition, and motor coordination are governed by widespread, intricate networks that span both cortical and subcortical areas. Nonhuman primate models are indispensable for advancing our understanding of the evolution of these networks and provide unique opportunities for experimental interventions that are not feasible in humans. In this study, we utilized movie-driven fMRI (md-fMRI) to investigate and delineate homologous functional networks in the common marmoset (*Callithrix jacchus*). Both marmosets and human subjects watched the same movie which incorporated a variety of visual and auditory stimuli. This method enabled the identification of potential homologues of large-scale functional networks involved in visual, auditory, cognitive, motor, and limbic functions in marmosets, offering new insights into the shared neurofunctional architecture across species.

## Introduction

The common marmoset (*Callithrix jacchus*) is a New-World monkey garnering increasing interest in neuroscience(1). This small non-human primate, characterized by rapid sexual maturity and high reproductive potential, shares significant neural and cognitive architecture similarities with both Old-World monkeys and humans. Notably, these similarities include a granular and complex prefrontal cortex(2), a sophisticated visual system(3), and comparable visuo-motor behavior(4,5). Recent studies have further revealed an extensive network dedicated to processing conspecific vocalizations, consistent with the high degree of social communication typical of these animals(6).

With the rising focus on marmosets, it becomes crucial to explore the cognitive-functional architecture of their brains, highlighting both homologies and differences compared to the human brain. This investigation, the primary aim of the present study, lays the groundwork for an interpretative framework for future comparative research involving this New World monkey. To achieve this goal, functional MRI (fMRI) is a pivotal tool, capable of capturing the hemodynamic response of the entire brain through a diverse array of experimental tasks (task-based fMRI) or resting-state paradigms (resting-state fMRI, or RS-fMRI). Both methodologies have already proven successful in comparative studies involving marmosets and humans, either by comparing brain activations in response to identical stimulation(7,8) or by identifying homologous large-scale brain networks between the two species during resting-state periods(9–11).

Despite their significance in understanding brain functioning, both task-based and RS-fMRI have some limitations. RS-fMRI, for instance, can capture and correlate the activity timecourse from multiple brain areas, investigating the functional connectivity of different “resting” regions. However, considering brain regions in isolation overlooks their interplay within broader functional networks(12,13), in which different areas perform different tasks to produce behaviorally relevant information. Furthermore, some "blocks" of these networks may only be active in the presence of certain stimuli or tasks, and thus RS-fMRI may not effectively and finely segregate entire networks. Finally, because resting-state patterns are state-agnostic and spontaneous, this technique cannot map the functional correspondences of interspecies blood oxygen level–dependent (BOLD) fluctuations over time.

On the other hand, task-based fMRI paradigms, also known as "functional localizers", can map specific functional networks using diverse stimuli. This approach allows for the comparison of BOLD fluctuations between different species in response to the same stimuli and has been widely applied in studies involving humans^3^, macaques(14,15), and marmosets(16–21). However, mapping the entire brain cortex of a primate would require numerous localizers, making this methodology impractical. Additionally, the repetition of the same stimulus over long periods, necessary to finely map functional networks, often faces challenges due to reduced compliance from non-human primates.

To address these limitations, movie-driven fMRI (md-fMRI), has emerged as a promising solution. In md-fMRI, naturalistic movies offer a continuously changing source of visual and auditory stimulation, maintaining the interest of non-human primates during fMRI sessions and creating a more ecologically valid stimulation condition compared to classic functional localizers. Studies in humans(22–25), macaques(26,27) and marmosets(28) have demonstrated that naturalistic movies recruit a wide range of brain areas, inducing highly selective responses between brain regions and highly reliable responses between subjects.

In this study, we presented the same movie, featuring a variety of visual (marmosets, humans, macaques, etc.) and auditory (marmoset vocalizations, speech, natural sounds, etc.) stimuli, to 8 common marmosets and 19 healthy humans. This approach allowed us to extract the average BOLD activation timecourses of 13 well-characterized functional networks from our human sample and correlate them with the voxel-by-voxel timecourses of our marmoset sample. These large-scale functional networks span different domains: visual (such as the action observation and face processing networks), auditory (vocalization processing and language comprehension network), cognitive (default mode and theory of mind network), motor (reaching/grasping and saccade network), and limbic (fear network). Our results reveal large-scale networks in marmosets that are putative functional homologues of various human brain networks.

## Methods

### Common marmosets

All experimental methods described were performed in accordance with the guidelines of the Canadian Council of Animal Care policy and a protocol approved by the Animal Care Committee of the University of Western Ontario Council on Animal Care. Animals were monitored during the fMRI acquisition sessions by a veterinary technician.

Eight adult marmosets (4 females, 28-45 months, mean age: 39.8 months) were the subjects in this study. For the head-fixed experiments, each animal underwent a surgery for the implantation of a head post(29), as detailed in our recent method article(30). Briefly, the procedure involved placing the animals in a stereotactic frame (Narishige, model SR-6C-HT) under gas anaesthesia (0.5-3% isoflurane). After making a midline incision along the skull, the skull surface was prepared with adhesive resin (All-Bond Universal; Bisco, Schaumburg, IL), and a PEEK head post was affixed using a resin composite (Core-Flo DC Lite; Bisco). Throughout the surgery, vital signs including heart rate, oxygen saturation, and body temperature were continuously monitored. Following a two-week recovery period, the marmosets underwent a three-week acclimatization procedure to the head-fixation system in a mock MRI environment. Detailed procedures can be found in (30).

### Human participants

Nineteen healthy humans (11 females, aged 25-45 years, mean age: 32.7 years) participated in the experiment. All participants were right-handed, reported normal or corrected-to-normal vision, and had no history of neurological or psychiatric disorders. Fourteen participants had prior experience with fMRI experiments. Prior to participation, subjects were fully informed about the experimental procedures and provided written consent. The study protocol was approved by the Human Ethics Committee of the University of Western Ontario.

### Stimuli

Both human participants and marmosets were presented with a movie that included audio. This movie was created by alternating baseline periods (a black circle of 0.36 visual degrees placed at the center of a grey screen) with excerpts from two different nature documentaries. The first, narrated in Spanish, is the documentary "Monkey Kingdom" produced by Disneynature in 2015; the second, narrated in English, is the episode "Urban Jungles" from the series "Hidden Kingdoms" released by the British Broadcasting Corporation in 2014. In the selected excerpts from these documentaries, a wide variety of visual stimuli (marmosets, humans, macaques, other animals, natural landscapes and city scenes, daytime and nighttime scenes, objects, food) and auditory stimuli (music, English/Spanish narration, marmoset and macaque vocalizations, natural and artificial sounds) were present. During the baseline periods, no auditory or visual stimuli (except for the central black circle used to attenuate the magnetic field-induced nystagmus) were presented. The movie’s total duration was 33 minutes, of which 5 minutes and 50 seconds were baseline periods. The remaining time featured excerpts from the two documentaries.

### Experimental Setup

During the scanning sessions, the marmosets were seated in a sphinx position in a custom-designed 3D-printed chair within the scanner. Their head was fixed using a head fixation system, securing the surgically implanted head post to a clamping bar (see (29,30) for details), and the two halves of the coil housing were positioned on either side of the head. After the head fixation step, the MRI-compatible auditory tubes (S14, Sensimetrics, Gloucester, MA) were positioned within the ear canals of the animal and kept in position using reusable sound-attenuating silicon earplugs (Amazon) and self-adhesive veterinary bandage (for all details, see (30)). An MRI-compatible camera (Model 12M-I, MRC Systems GmbH) mounted on the frontal part of the chair allowed to monitor the animal’s activity. However, the recordings’ quality was insufficient for detailed eye-tracking analysis, due to the challenge of tracking the large marmoset’s pupils once the eyelids are not fully open, as it often happened in the 33-minutes long functional runs. Within the scanner, monkeys faced a translucent screen placed 119 cm from their eyes, onto which visual stimuli were projected using an LCSD projector (Model VLP-FE40, Sony Corporation, Tokyo, Japan) via back-reflection on a first surface mirror. Visual stimuli were presented using PowerPoint software (version 16.83, Microsoft Corporation, WA, USA) and synchronized with MRI TTL pulses triggered by a Raspberry Pi (model 3B+, Raspberry Pi Foundation, Cambridge, UK) operating via a custom-written Python program. During scanning sessions, the monkeys were given a liquid reward consisting of a mixture of melted marshmallows in water. A plastic tube was positioned on the animal’s lips, delivering a single drop of the reward every 3 TRs to facilitate monkey’s vigilance. Each marmoset was presented with the same movie five times in five separate fMRI sessions.

Human subjects lied supine in the MR scanner and viewed stimuli via a rear projection system (Avotech SV-6011, Avotec Incorporated) through a surface mirror attached to the head coil. As for marmosets, visual stimuli were presented using PowerPoint software and synchronized with MRI TTL pulses triggered by the Raspberry Pi via a custom-written Python program. Before positioning the head in the coil, participants were asked to insert the MRI-compatible auditory tubes (T14, Sensimetrics, Gloucester, MA) in their ear canals, and were asked to confirm that the volume of the auditory stimuli was appropriate. Human participants watched the movie only once.

### MRI data acquisition

Both marmoset and human data were collected at the Center for Functional and Metabolic Mapping at the University of Western Ontario.

Marmoset data were acquired on a 9.4T 31-cm horizontal bore MR scanner (Varian) with a Bruker BioSpec Avance III HD console running the software package Paravision-360 (Bruker BioSpin Corp) using a custom-built high-performance 15-cm diameter gradient coil (maximum gradient strength: 1.5 mT/m/A), and a custom-built eight-channel receive coil. Preamplifiers were located behind the animals, and the receive coil was placed inside an in-house built quadrature birdcage coil (12-cm inner diameter) used for transmission. We acquired 5 functional runs for each animal in 5 separate sessions using gradient-echo based single-shot echo-planar images (EPI) sequence with the following parameters: TR=1.5s, TE = 15ms, flip angle = 40°, field of view=64x48 mm, matrix size = 96x128, resolution of 0.5 mm^3^ isotropic, number of slices= 42 [axial], bandwidth=400 kHz, GRAPPA acceleration factor: 2 (left-right). Moreover, a second set of EPIs with an opposite phase-encoding direction (right-left) was collected to correct for the EPI-distortion. Finally, a T2-weighted structural image was also acquired for each animal during one of the sessions with the following parameters: TR=7s, TE=52ms, field of view=51.2x51.2 mm, resolution of 0.133x0.133x0.5 mm, number of slices= 45 [axial], bandwidth=50 kHz, GRAPPA acceleration factor: 2.

For human subjects, fMRI data were acquired on a 7T 68-cm MRI scanner (Siemens Magnetom 7T MRI Plus) with an AC-84 Mark II gradient coil, an in-house built 8-channel parallel transmit, and a 32-channel receive coil(31). Each participant underwent a single functional run using Multi-Band EPI BOLD sequences with the following parameters: TR=1.5s, TE = 20ms, flip angle = 30°, field of view=208x208 mm, matrix size = 104x104, resolution of 2 mm^3^ isotropic, number of slices= 62, GRAPPA acceleration factor: 3 (anterior-posterior), multi-band acceleration factor: 2. Field map images were also computed from the magnitude image and the two phase images. An MP2RAGE structural image was also acquired for each subject with the following parameters: TR=6s, TE=2.13 ms, TI1 / TI2 = 800 / 2700 ms, field of view=240x240 mm, matrix size= 320x320, resolution of 0.75 mm^3^ isotropic, number of slices= 45, GRAPPA acceleration factor (anterior posterior): 3.

### fMRI data preprocessing

For marmoset fMRI data, preprocessing utilized functions from AFNI(32) and FSL(33) software packages. Raw DICOM images were converted to NIfTI format using dcm2niix (AFNI) and reoriented with fslswapdim and fslorient (FSL) functions. Given a TR of 1.5 seconds, each functional run comprised 1325 full volumes. Prior to preprocessing, the first and last 5 volumes were discarded (using AFNI’s 3dcalc function) due to altered signal (both the first and last 5 volumes corresponded to baseline periods in the movie), resulting in 1315 final volumes. Distortion correction was applied using FSL’s topup and applytopup functions. Following this step, data underwent despiking (3dDespike, AFNI), time-shifting (3dTshift, AFNI), and registration to the base volume (the middle one for each run) using 3dvolreg (AFNI). Spatial smoothing with a 1.5 mm full-width half-maximum (FWHM) Gaussian kernel was performed using 3dmerge (AFNI), followed by temporal filtering (0.01-0.5 Hz) with 3dBandpass (AFNI), and normalization using 3dDetrend (AFNI). Mean functional images were calculated for each run and linearly registered to the corresponding animal’s anatomical image -manually skull-stripped using FSL eyes tool- using FLIRT (FMRIB’s Linear Image Registration Tool). The resulting transformation matrix was then applied to register all functional volumes to the animal’s anatomical image, followed by masking to remove non-brain voxels. Finally, the anatomical image was linearly registered to the NIH marmoset brain template(34) using ANTs (Advanced Normalization Tools)(35), and the resulting transformation matrix was applied to register functional volumes to the same template.

For human fMRI data, preprocessing was performed primarily with SPM12 (Statistical Parametric Mapping, Wellcome Center for Human Neuroimaging, London, UK) following conversion from DICOM to NIfTI and the removal of the first and last 5 volumes (AFNI’s dcm2nixx and 3dcalc functions respectively). Functional images underwent realignment for motion correction and slice-timing correction. Field map correction was applied to each run using magnitude and phase images. The functional images were then coregistered with each participant’s MP2RAGE structural scan and subsequently normalized to the Montreal Neurological Institute (MNI) standard brain space. Anatomical images were segmented into gray matter, white matter, and CSF partitions and normalized to MNI space. Functional images were finally smoothed using a FWHM Gaussian kernel three times larger than the voxel size (6 mm isotropic), temporally filtered (0.01-0.5 Hz) using 3dBandpass (AFNI), and normalized using 3dDetrend (AFNI), mirroring the preprocessing steps applied to the marmoset data.

### Functional networks selection

The main objective of this study was to identify functional homologies of human long-range brain networks in marmosets. Considering the diverse range of stimuli presented in the movie, we curated a selection of 13 functional networks from the existing human fMRI literature. These networks cover key domains including visual and auditory processing, motor control, cognition/executive functions, and emotion. Subsequently, the average timecourses of these networks, derived from our human sample, were correlated with the timecourse of each voxel in the marmoset brain. This analysis aimed to identify regions within the marmoset brain exhibiting similar functional activation patterns (for detailed methodology, refer to the Statistical Analysis section). Additionally, correlation maps were compared, where available, with group activation maps derived from previous task-based studies in marmosets conducted by our group(6,11,16,20,21). Supplementary Figure 1 illustrates the extracted human networks, their respective sources, and, when available, the corresponding marmoset task-based network with its source information. The masks of the human networks and the functional maps from marmoset studies are presented in the Results section (Figures 2 to 7).

All network masks were resampled using AFNI’s 3dresample function to achieve a resolution matching to our human fMRI data (2 mm isotropic) and were aligned to the MNI standard brain space. Marmoset functional maps, which already matched the resolution of our fMRI data (0.5 mm isotropic), were registered to the NIH marmoset brain template(34).

### Statistical Analysis

To generate a consolidated representation for each species and voxel, we conducted separate averaging procedures on the normalized timecourses of each voxel across our human and marmoset samples. Utilizing a custom-written MATLAB (R2021b, The MathWorks Inc.) script, we derived a single 4D fMRI map for each voxel for both the human and marmoset samples. These two 4D fMRI maps represented the average timecourse (1 to 1315 volumes) of each voxel across 19 (humans) and 40 (8 marmosets, 5 runs per marmoset) functional runs, respectively.

To assess the validity of our approach, we initially extracted the timecourse of area MT, a well-defined motion-processing brain region in both humans and marmosets. This region should thus respond well to the movie in both humans and marmosets. To validate this assumption, we extracted the timecourse of the MT area delineated by the multi-modal cortical parcellation atlas(36) in humans and by the Paxinos(37) parcellation of the NIH marmoset brain atlas(34). Firstly, we computed the within-species functional connectivity correlating the human and marmoset MT timecourse to all the voxels in the human (Figure 1C and 1E) and marmoset brain (Figure 1D and 1F), respectively. These maps show reliable correlations of the MT area with several other visual, temporal and parietal regions in both species. Importantly, as depicted in Figure 1, the timecourses of these two regions also exhibited a robust correlation between-species (r = 0.49), confirming the reliability of our approach. To further verify this, we correlated the timecourse of human MT to all the voxels in the marmoset brain (Figure 1G). This between-species correlation revealed a functional connectivity map sharing important overlap with the within-species marmoset correlation map (Figure 1D), supporting a functional homology between human and marmoset MT areas.

**Figure 1.**
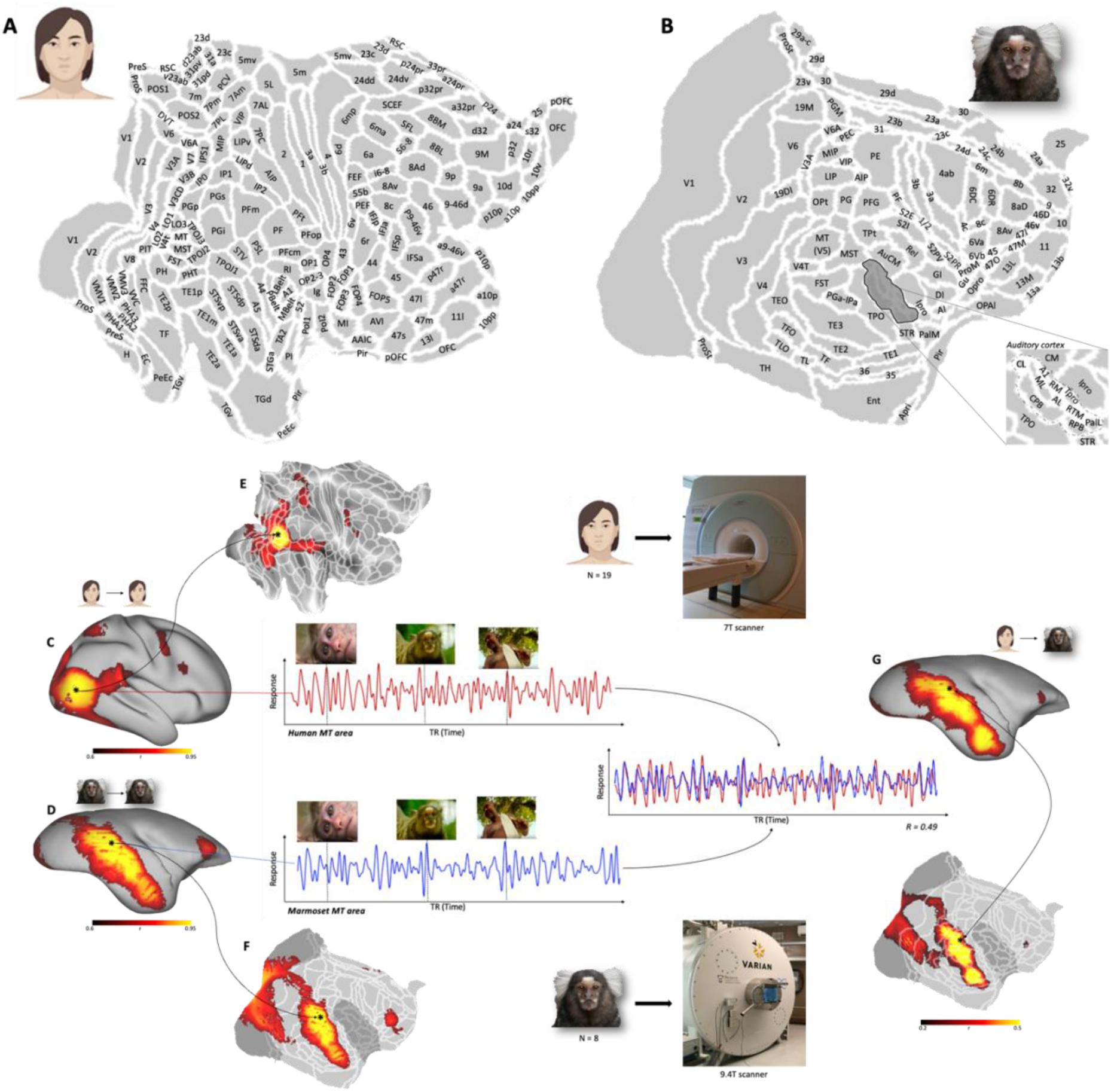
BOLD timecourse correlation between human and marmoset. In panels A and B, the representation of the human (A) and marmoset (B) brains as flat maps. White lines delineate the multi-modal cortical parcellation atlas(36) and the Paxinos parcellation(37) of the NIH marmoset brain atlas(34), for humans and marmosets respectively. At the center and at the bottom of the figure, the 7 Tesla (center) and 9.4 Tesla (bottom) scanners utilized for our fMRI sessions on healthy humans (n = 19) and common marmosets (n = 8, 5 functional runs per marmoset), respectively. On the left side of the figure, right lateral fiducial surfaces of the human (C) and marmoset (D) brain display the localization of motion-selective area MT in the two species. On both surfaces is displayed the map resulting from the within-species correlation of the timecourse of the MT area with each voxel of the brain. The same correlation maps are represented on flat maps for both humans (E) and marmosets (F). White lines delineate the multi-modal cortical parcellation atlas(36) (E) and the Paxinos parcellation(37) of the NIH marmoset brain atlas(34) (F). The central graphs show the average timecourse of the human (red) and marmoset (blue) area MT during the presentation of the 33-minutes long naturalistic movie. The pictures of a macaque, a marmoset and a human are real frames taken from the experimental movie, but their temporal position on the graph does not reflect their real timing. The correlation between these two timecourses (r = 0.49) highlight the reliability of our method. Panel G shows the correlation map of the human area MT timecourse with all marmoset cortical voxels on the right lateral fiducial surface and flat map (white lines delineate the Paxinos parcellation(37) of the NIH marmoset brain atlas(34)). Marmoset MT area is highlighted in both maps with a black asterisk.

Subsequently, we utilized this approach and extracted the timecourses corresponding to each of the selected human networks. Given that meta-analytic maps from Neurosynth often contain numerous isolated single voxels dispersed throughout the brain, we binarized each human network map and subsequently removed isolated voxels using AFNI’s 3dcalc function with the *-isola* option. Following this first step, we employed a custom MATLAB script to extract and average the timecourses of all voxels encompassed within each network mask, resulting in 13 distinct human timecourses. Finally, we conducted pairwise correlations between these timecourses and I) the timecourse of each voxel in the human brain, and II) the timecourse of each voxel in the marmoset brain. To ensure specificity in our analysis, we restricted correlations to voxels within the cortical Paxinos(37) parcellation of the NIH marmoset brain atlas(34) and, subcortically, within the recently published Subcortical Atlas of the Marmoset (SAM)(38). For the human brain, analyses were restricted to voxels within the multi-modal cortical parcellation atlas(36).

As the final step, we investigated the anatomical overlap of the functional networks we identified through within-species and between-species correlations. Our goal was to identify potential multifunctional hubs, which are brain regions involved in multiple functional networks in both species. It’s important to note that the networks described here are selected examples covering different cognitive domains. Different results may emerge with the selection of different networks, masks of the described networks, or modifications to the statistical thresholds for more or less permissive functional homologies.

To create a coincidence map, we transformed the correlation maps into binary masks. Voxels with values above a specific statistical threshold were coded as "1," while all other voxels were coded as "0." For the within-species correlation maps (correlating the timecourse extracted from each functional network mask with every voxel of the human brain), we used a threshold of r>0.6. For the between-species correlation maps (correlating the timecourse of each functional network mask with every voxel of the marmoset brain), the thresholds varied by network type: r>0.25 for perceptual networks (visual, auditory, and multisensory) and the speech production network, r>0.2 for the fear network, and r>0.15 for cognitive and motor networks.

Finally, we computed the coincidence map by summing the 13 binary masks and dividing the final map by the number of selected networks (13). This process yielded a coincidence map with values ranging from 0 (voxel never included in the correlation maps) to 1 (voxel above threshold in all correlation maps).

## Results

### Visual networks

We first tried to identify marmoset functional homologues of human long-range networks involved in visual processing, including the action observation network (AON), the face processing network, and the place network. For brevity, the results discussed in the main text pertain solely to the right hemisphere of both marmosets and humans.

The human AON is a well-established long-range network comprising areas activated during the observation of goal-directed actions(39–41). As depicted in Figure 2A, this network encompasses visual (V4, V4t, V7, MT), temporal (MST, FST, temporo-parieto-occipital junction -TPOJ-, PHT, superior temporal visual area -STV-, PF complex), parietal (LIP, AIP, 7PC), and premotor areas (6mp, 6a, 6d, 6v, FEF). The correlation of the timecourse of this network mask with all the voxels in the human brain produced a network highly overlapping with the mask, with further inclusion of some high-level visual areas. This difference was probably driven by the experimental material we adopted, rich with a wide variety of moving visual stimuli. Notably, the timecourse of this human network mask exhibits robust correlations with a number of areas in the marmoset brain, including visual areas such as V1, V2, V3, V4, V4T, and MT, together with temporal regions MST, FST, TPO, PGa-IPa, and prefrontal cortical regions including areas 45 and 8Av. Moreover, we observed correlations with the human AON timecourse subcortically, in the pulvinar, putamen, caudate and in the lateral geniculate nucleus (LGN). This activation pattern resembles the human AON network. It also closely resembles our previous findings in which marmosets observed goal-directed or non-goal-directed actions(20). The marmoset task-based AON, as depicted in Figure 2A, includes an occipito-temporal cluster involving V2, V3, V4, V4t, MT, MST, FST, Pga-IPa, and TE3, along with a prefrontal cluster encompassing areas 45, 8Av, and additional areas within the dorsal and ventral portions of area 6, area 47, and area 8C. Furthermore, the medial wall of the right hemisphere reveals the inclusion of a portion of the posterior cingulate cortex, including areas 23v, 29a-c, 29d, 30, and PGM. Subcortically, the observation of goal-directed actions recruited the pulvinar, the ventral lateral and ventral posterior nuclei of the thalamus and the superior colliculus (SC).

**Figure 2.**
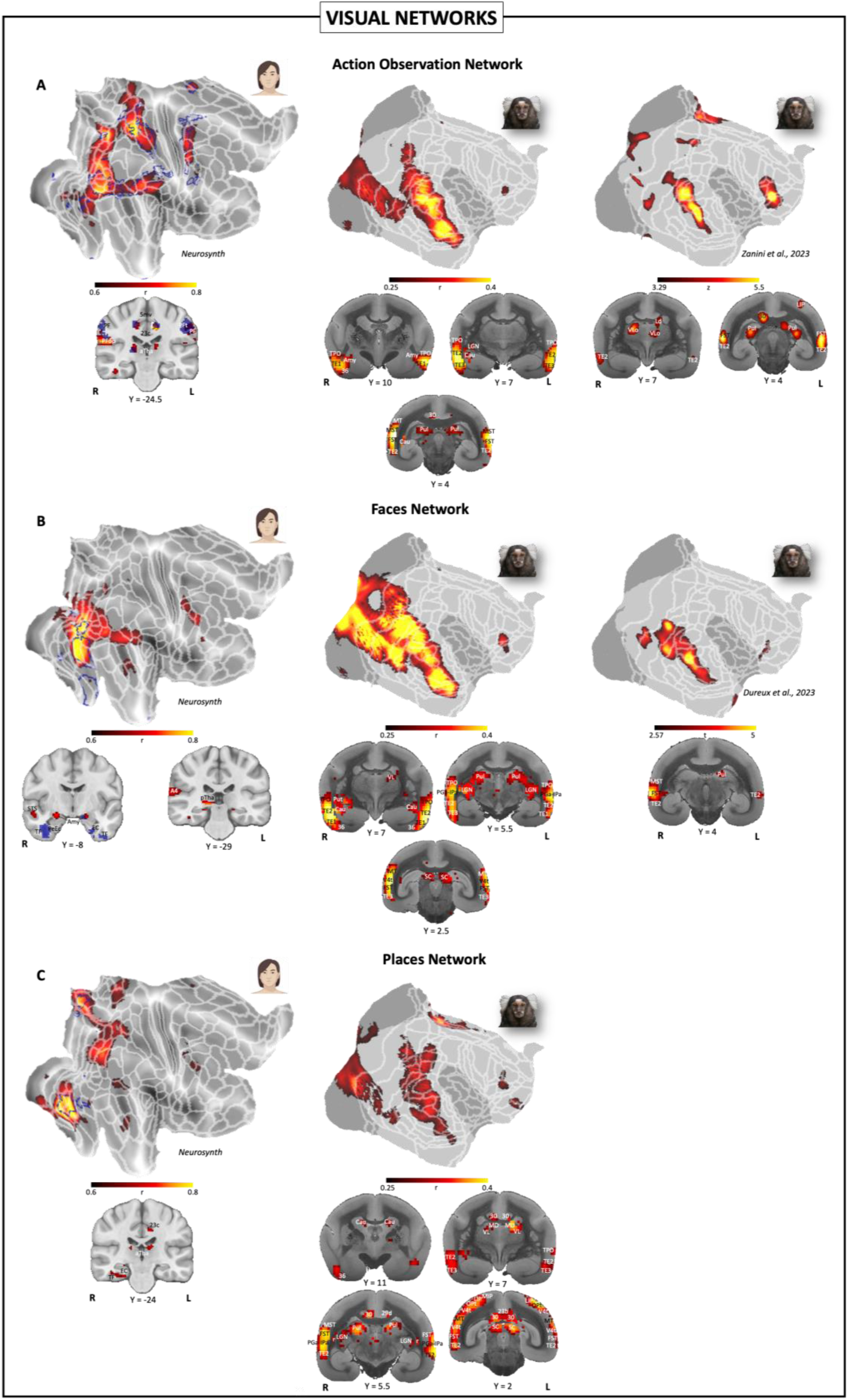
Visual functional network correlation between human and marmosets. The figure displays the human-marmoset correlations concerning A) the action observation network, B) the face processing network, and C) the place network. In the left column of each panel, the masks used to extract the network timecourse from our human sample (blue colour) are displayed on flat maps of the right hemisphere. Hot colour represents the within-species correlation of the timecourse of the network with all the voxels of the brain. White lines delineate the multi-modal cortical parcellation atlas(36). In the central column, the correlation maps obtained correlating the human network timecourses to the marmoset brain timecourse, on a voxel-by-voxel basis. In the right column, when available, the group activation maps obtained by our research group in previous studies (from top to bottom: goal-directed versus non-goal-directed actions(20), static faces vs static objects(16)) on common marmosets. Both correlation and group activation maps are displayed on flat maps of the right hemisphere. White lines delineate the Paxinos parcellation(37) of the NIH marmoset brain atlas(34). For all maps, the volumetric cortical and subcortical activations at different inter-aural levels are displayed on the coronal slices of MR anatomical images of a human (left column) or a marmoset (central and right columns) brain.

The second visual network that we investigated is the network activated by the perception of faces(42–44). This face-processing network encompasses the ventral portions of the occipital and temporal lobes in humans, including V4t, PH, the ventral visual complex (VVC), the posterior inferotemporal area (PIT), the ventral portion of area TE2 and the fusiform face complex (FFC). Further activations are observed in the perirhinal ectorhinal cortex (PeEc) and, subcortically in the amygdala. The voxel-by-voxel correlation of the timecourse of this network mask revealed an extensive overlapping of the aforementioned areas, but also the involvement of many additional regions, in particular at the occipito-temporal and premotor levels. This difference, as already mentioned for the AON, could be due to the fact that the faces in our movie were not presented in an isolated and static fashion, but in motion and in the presence of numerous other stimuli. As depicted in Figure 2B, the timecourse extracted from this network in our human sample demonstrates robust correlation with a broad occipito-temporal pattern of areas in the marmoset brain, spanning visual areas V1, V2, V3, V4, V4t, MT, and 19DI, as well as temporal regions MST, FST, PGa-IPa, TPO, TEO, the TE complex, and ventral temporal areas 36 and 35. Moreover, reliable correlations extended into the prefrontal lobe, involving areas 8Av, 45, and 47. Subcortically, the network includes SC, LGN, pulvinar, putamen, caudate and the lateral and basal nuclei of the amygdala. Although this network appears to be extensive in the marmoset brain compared to humans, evidence supporting this correlation map stems from another study conducted by our research group(16). Dureux and colleagues utilized a localizer task comparing brain activations induced by the presentation of static images of faces and objects. The face processing network identified in that study (and also earlier (17,18)) is highly coherent with the correlation map obtained through md-fMRI here, involving a similar occipito-temporal and a prefrontal cortical cluster, plus the basal nucleus of the amygdala subcortically. Additional activations are observed in the orbital proisocortex (Opro) and, subcortically, in the claustrum. All these results are represented in Figure 2.

Lastly, we aimed to identify a functional equivalent in the marmoset for the third human visual network, known as the scene (or ‘place’) network. This network consists of regions that process "places," defined as real-world environments featuring background elements and various distinct stimuli such as objects, people, and animals(45,46). The mask we used to extract the timecourse of this network includes a cluster among the parahippocampal areas (PHA1, PHA2, and PHA3) and the ventromedial visual areas (VMV1, VMV2, and VMV3) in the occipital lobe, along with a smaller cluster bridging the FFC and the TE2p area. Additionally, an activation spot at the occipito-parietal junction covers the parieto-occipital sulcus area 1 (POS1) and the dorsal transitional visual area (DVT). These clusters, except for the spot in FFC, are found in the map obtained through within-species correlation of the timecourse of this mask with all voxels of the human brain. Moreover, robust correlations are observed in the retrosplenial complex (RSC), posterior cingulate cortex (5mv, 23c, and precuneus), and the occipito-parietal region, including V3B, V3CD, PGp, IP0, IPS1, and the intraparietal sulcus areas MIP, LIPv, and LIPd. Lower correlations are seen in the temporal (STS, TPOJ1) and frontal (IFJa and IFJb) regions. At the subcortical level, a strong correlation is observed with the anterior thalamus. The correlation of the timecourse of the human places network with the timecourse of the marmoset brain reveals two robust correlation spots. The first, located in the posterior cingulate cortex, includes areas 30 and 29d. The second is situated in the occipital lobe, encompassing portions of areas V1, V2, V3, and V4. Besides the posterior cingulate cortex, another parallel between the human and marmoset places networks is found at the occipito-parietal junction (19DI, V3A, MIP, LIP, and Opt). In the marmoset, this correlation cluster extends further into the temporal lobe, covering areas MT, V4t, MST, FST, PGa-IPa, and TE3. Cortical correlations include areas 8Av, 8aD, and 6DR and a second orbitofrontal spot (bridging areas 13 and 11). Subcortical correlations are more extensive than in the human map, including the caudate nucleus, multiple thalamic regions (mediodorsal nucleus or MD, ventral lateral nucleus or VL, pulvinar, and lateral geniculate nucleus or LGN), and the superior colliculus. All these results are depicted in Figure 2C.

### Auditory networks

Next, we tried to identify functional homologues of human networks involved in different aspects of auditory processing. For brevity and considering the lateralization of the language networks in the human brain, the results represented and discussed in the main text focused only on the left hemisphere of both humans and marmosets.

We first attempted to identify a marmoset homologue of the human vocalization network, which is activated by vocal sounds. This network (from (47)) includes mainly areas located along the superior temporal sulcus (TPOJ, STV, STS and STGa) and the auditory cortex (A1, A4, A5, Pbelt, Lbelt, Mbelt and RI). More anteriorly, the network includes a cluster spanning the insular and frontal cortices (the frontal opercular area or FOP, OP4, 6r, 43, and 44) and two smaller regions, one coinciding with area 55b and the other with the superior frontal language area (SFL). The correlation of the timecourse of this mask with the human brain shows robust correlations only with the more posterior cluster, located in the temporal and auditory cortices. The more anterior clusters emerge only at lower levels of correlation (not presented in the current manuscript). Correlating the timecourse of this human network to our marmoset data revealed a pattern of areas spanning auditory cortices and the superior temporal lobe. In particular, the correlation map included auditory areas A1, CL, CM, ML, RM, RTL, RPB and CPB, the insular cortex (DI, GI, Ipro, PalL), the temporal areas PGa-IPa, MST, FST, TPO and the conjunction between areas TE1, TE2, TE3 and 36. Furthermore, correlated clusters are observed in the occipital lobe, and on the medial wall in cingulate area 32. Subcortically, we found correlations in the superior and inferior colliculus, the PAG, the globus pallidus, the pulvinar and the centromedian and mediodorsal nuclei of the thalamus. Recently, we investigated the neural activations evoked by the presentation of conspecific calls in common marmosets(6). The group activation map, illustrated in Figure 3A, shows similar activations as the correlation map, with a more pronounced somatomotor component and the recruitment of a larger subcortical network.

**Figure 3.**
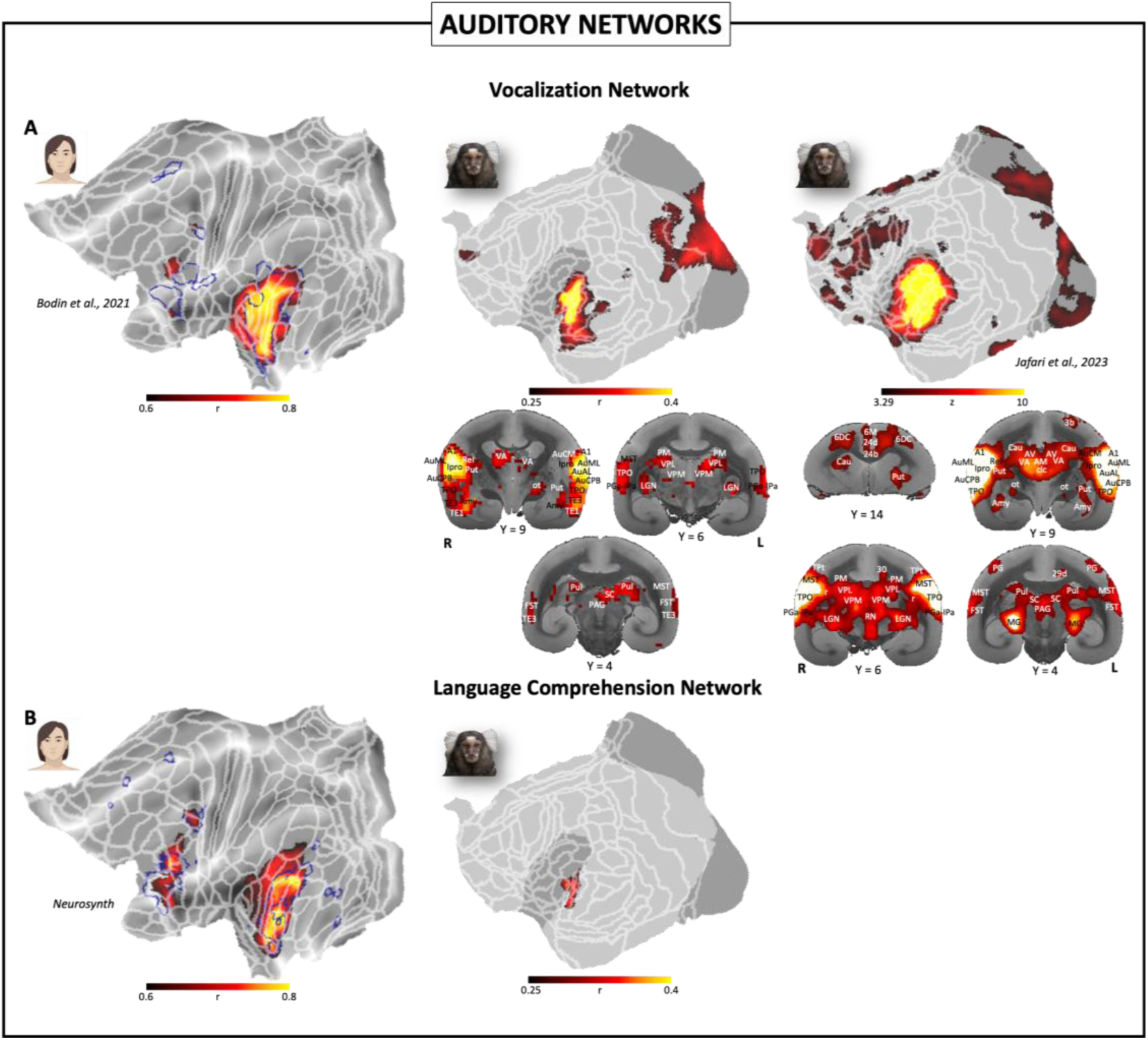
Auditory functional network correlation between human and marmosets. The figure displays the human-marmoset correlations concerning A) the vocalization network, and B) the language comprehension network. In the left column of each panel, the masks used to extract the network timecourse from our human sample (blue colour) are displayed on flat maps of the left hemisphere. Hot colour represents the within-species correlation of the timecourse of the network with all the voxels of the brain. White lines delineate the multi-modal cortical parcellation atlas(36). In the central column, the correlation maps obtained correlating the human network timecourses to the marmoset brain timecourse, on a voxel-by-voxel basis. In the right column, when available, the group activation maps obtained by our research group in previous studies (for the vocalization network: marmoset vocalizations versus baseline(6)) on common marmosets. Both correlation and group activation maps are displayed on flat maps of the left hemisphere. White lines delineate the Paxinos parcellation(37) of the NIH marmoset brain atlas(34). For the marmoset vocalization maps, the volumetric cortical and subcortical activations at different inter-aural levels are displayed on the coronal slices of MR anatomical images a marmoset (central and right columns) brain.

Next, we identified a putative marmoset homologue of the human language comprehension network. As shown in Figure 3B, the mask of this network included a cluster of activation extending across the STS and the auditory cortices A4 and A5, a second cluster in the precentral gyrus covering area 55b, the frontal (FEF) and the premotor (PEF) eye field, and a third cluster including the areas 6r, 44, 45 and 47l. Medially, the network recruited the SCEF, the superior frontal language area (SFL), and areas 8BM and 9m. The within-species correlation reported a pattern of results highly coherent with the mask described and showing further robust correlation with regions in the auditory and temporal cortices (as the perysilvian language area or PSL and the retroinsular cortex or RI). The correlation of this network timecourse with the marmoset timecourse showed a single cluster of correlations, centered on the auditory cortex. At a lower correlation threshold (data not shown here), this correlation cluster extends, and two new clusters appear—one in the visual cortex and a second in the anterior cingulate cortex (particularly area 32), making this putative marmoset network very similar to the one described for vocalization processing.

### Multisensory network

Staying within the domain of perceptual functional networks, we investigated the functional homologue in the marmoset of a human multisensory network, specifically the audiovisual integration network, responsible for integrating visual and auditory signals(48). This network encompasses several brain regions in humans including the superior temporal sulcus (STS), the TPOJ, and the STV. In the auditory cortex, the network comprises regions A1, A4 and A5, the lateral, medial and posterior belt complex (Lbelt, Mbelt, Pbelt), the perisylvian language area (PSL), the frontal opercular (FOP), the anterior ventral (AVI), the middle (MI) and the retroinsular (RI) areas. Finally, in the frontal lobe, the inferior frontal gyrus and the rostral area 6. Subcortically, the network mask included also a portion of the posterior thalamus. The within-species correlation confirmed this network, with robust correlations also extending to further portions of the STS and the auditory cortex (see Figure 4). Subcortically, a small correlation spot in the amygdala complements the confirmed presence of the posterior thalamus. Upon correlating the timecourse extracted from this network in our human sample with the marmoset brain’s timecourse, we observe correlations across a network of areas along the occipito-temporal axis, encompassing wide portions of the auditory cortex. Notably, this correlation map shows a reduced recruitment of the TE complex and dorsal visual areas, while incorporating several regions of the auditory cortex, including the core (primary area [A1] and rostral field [R], rostral temporal [RT]), belt (caudomedial [CM], caudolateral [CL], mediolateral [ML], rostromedial [RM], anterolateral [AL], rostrotemporal medial [RTM], rostrotemporal lateral [RTL]), and parabelt areas (caudal parabelt [CPB], rostral parabelt [RPB]). Further strong correlations are observed in the retroinsular (ReI), granular (GI) and dysgranular (DI) insular areas, in the medial and lateral parts of the parainsular cortex (PalM and PalL) and in the insular proisocortex (Ipro). Finally, reliable correlations are observed in the superior temporal rostral cortex (STR) and in temporoparietal transitional area (TPt). Subcortically, the audiovisual integration recruits the ventral lateral and anterior nuclei of the thalamus, the LGN, the SC, the pulvinar, the periaqueductal gray (PAG), the basal and lateral nuclei of the amygdala and the cerebellar lobule IX. We recently explored brain activations in the common marmoset elicited by the presentation of faces alone, marmoset vocalizations alone, or the combination of these two stimuli(21). This combined condition, compared to the baseline, offers a valuable insight into audiovisual integration in these New World monkeys. As illustrated in Figure 4, the audiovisual network recruited by this stimulus combination exhibits striking similarities to the correlation map described above. Notably, the occipito-temporal cluster demonstrates activations in visual areas such as V4, V4t, and MT, alongside engagements of areas MST, FST, PGa-IPa, and a prominent focus of activation at the conjunction of TE1, TE2, TE3, and TPO. Furthermore, activations in the auditory cortex are evident in core (A1), belt (CM, CL, RM, AL, RTM), and parabelt areas (CPB, RPB). Additional activations consistent with the correlation map derived from the movie task are observed in STR, TPt, and the insular cortex (GI, DI, PalL, PalM, Ipro, ReI), but with additional recruitment of the temporal (Tpro) and temporopolar (TPPro) proisocortex. Moreover, this multisensory network further includes the cingulate (29a-c, 29d, 30) and the parietal cortices (VIP, MIP, LIP, AIP, PG). Finally, a cluster of activation is evident in the prefrontal cortex, encompassing areas 45, 47, 8Av, 8aD, 8c and 6DR. Subcortically, activations consistent with the correlation map are observed in the pulvinar, the SC, the lateral nucleus of the amygdala, the anterior ventral nucleus of the thalamus, the centromedian and the central lateral thalamic nuclei.

**Figure 4.**
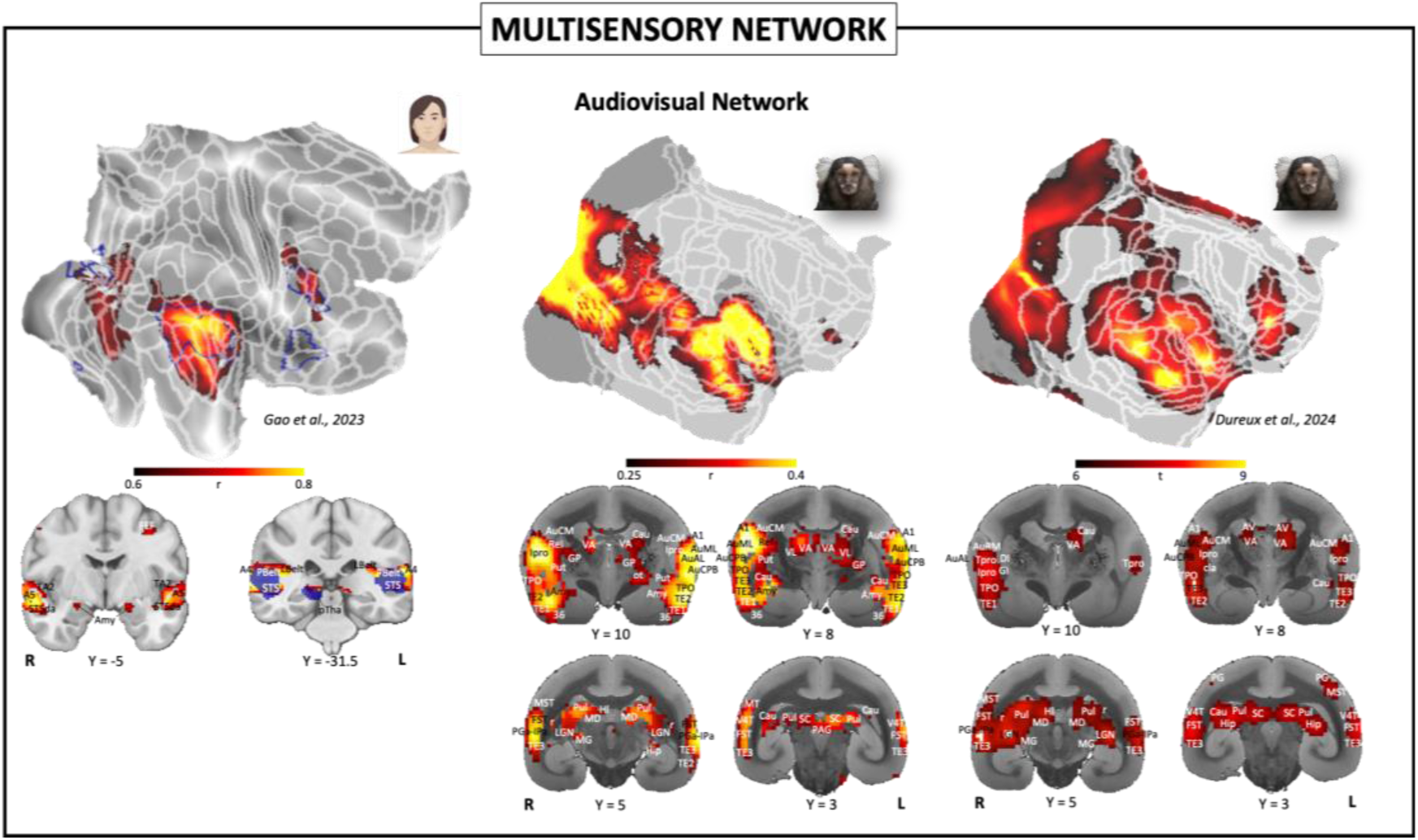
Audiovisual integration network correlation between human and marmosets. The figure displays the human-marmoset correlations concerning the audiovisual integration network. In the left column, the mask used to extract the network timecourse from our human sample (blue colour) is displayed on flat map of the right hemisphere. Hot colour represents the within-species correlation of the timecourse of the network with all the voxels of the brain. White lines delineate the multi-modal cortical parcellation atlas(36). In the central column, the correlation map obtained correlating the human network timecourse to the marmoset brain timecourse, on a voxel-by-voxel basis. In the right column, the group activation maps obtained by our research group in a previous study (marmoset vocalizations + marmoset faces versus baseline(21)) on common marmosets. Both correlation and group activation maps are displayed on flat maps of the left hemisphere. White lines delineate the Paxinos parcellation(37) of the NIH marmoset brain atlas(34). For the marmoset vocalization maps, the volumetric cortical and subcortical activations are displayed at different inter-aural levels on the coronal slices of MR anatomical images of a marmoset brain (central and right columns).

### Cognitive networks

In the cognitive domain, we considered several brain networks, including the default mode, the central executive and the theory of mind networks. Here, we focus on the right hemisphere for both human and marmoset brains.

We firstly focused on the human default mode network (DMN), which is defined as the pattern of areas more active during "rest" compared to task periods(49). In humans, this network shows three clusters of activation located in the PGi/PGs area, in the superior temporal sulcus (TE1) and in the frontal lobe (areas 8aD and s6-8) respectively. On the medial wall, the DMN includes a cluster of activations in the anterior cingulate and rostral prefrontal cortex (areas 32, 24, 10 and 9) and in the posterior cingulate cortex (areas 23ab, 23c, 23d, 31, RSC and 7m). Subcortically, the human DMN also encompasses the hippocampus. The within-species correlation of the timecourse of this functional network revealed a pattern where the most robust correlations are confined to the previously described mask, with some further lower correlations at the local level. Subcortically, correlations are observed in the anterior thalamus and the nucleus accumbens (NA). When we correlated the timecourse of this network to the timecourses of the voxels in the marmoset, we did not find correlations in the temporal cortex, but some correlations in the parietal lobe including MIP, LIP, PEC, PE and Opt. In the frontal cortex, the marmoset putative DMN includes the dorsal area 6, areas 8aD and 8C and orbitofrontal area 13. Notably, the medial wall correlation map strongly resembles the human DMN, with a cluster of correlation in the medial prefrontal cortex (areas 32, 8b and 24) and one in the posterior cingulate cortex (area 29a-c, 29d, 23a, 23b and PGM). A similar pattern of areas had been observed in a previous study of our research group through independent component analysis (ICA)(11) (Figure 5A). In fact, Hori and colleagues reported a network that included parietal, prefrontal and posterior cingulate components which strongly overlapped with the correlation map described here. Lowering the statistical threshold (data not shown here), the medial prefrontal cortex cluster is also present in the ICA map. Subcortically, the marmoset DMN identified by ICA also included the hippocampus.

**Figure 5.**
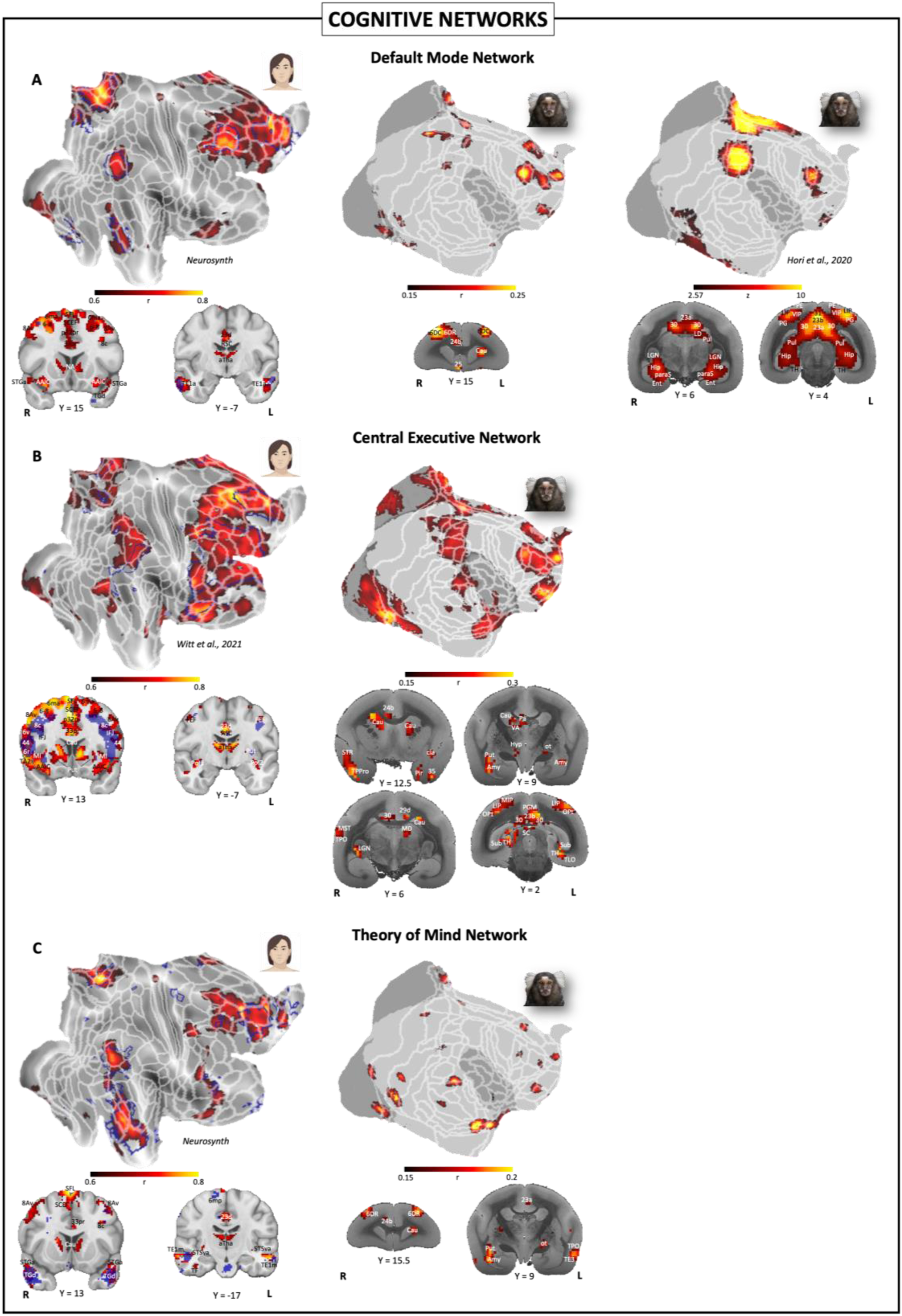
Cognitive functional network correlation between human and marmosets. The figure displays the human-marmoset correlations concerning A) the default mode network, B) the central executive network, and C) the theory of mind network. In the left column of each panel, the masks used to extract the network timecourse from our human sample (blue colour) are displayed on flat maps of the right hemisphere. Hot colour represents the within-species correlation of the timecourse of the network with all the voxels of the brain. White lines delineate the multi-modal cortical parcellation atlas(36). In the central column, the correlation maps obtained correlating the human network timecourses to the marmoset brain timecourse, on a voxel-by-voxel basis. In the right column, when available, the group activation maps obtained by our research group in previous studies (for the default mode network: independent component analysis on resting-state data(11)) on common marmosets. Both correlation and group activation maps are displayed on flat maps of the right hemisphere. White lines delineate the Paxinos parcellation(37) of the NIH marmoset brain atlas(34). For all maps, the volumetric cortical and subcortical activations at different inter-aural levels are displayed on the coronal slices of MR anatomical images of a human (left column) or a marmoset (central and right columns) brain.

The second cognitive network that we investigated was the central executive network(50). In humans, this network encompasses areas in the temporal lobe (STV, TPOJ, STS, PFm), the auditory cortex (A5, PSL), the insular cortex (MI, AVI, FOP) and a large portion of the frontal lobe, including areas 6r, 8aD, 8Av, 8C, 9a,10p, 44, 45, and 46, plus the IFS and IFJ. Moreover, clusters of activations are observed in the medial prefrontal cortex (areas 9m, 8BM, 24 and 32), in the posterior cingulate cortex (areas 31, RSC and 23), plus the parieto-occipital area POS2. The within-species correlation map revealed results coherent with this mask, with some further robust correlation on the medial wall (SCEF, 32pr, 33pr and 24pr in particular) and in the orbitofrontal cortex (areas 13 and 11) depicted in Figure 5B. Subcortically we found correlations with the anterior portion of the thalamus, the caudate and the putamen. On the other hand, the between-species correlation map of this network reveals a network recruiting parietal (MIP, LIP, PEC, PE, PG, Opt), temporal (MST, TPt, TPO, STR, TE1, 36, 35 and PalM), and prefrontal areas (dorsal area 6, 8, 11, 13 and 46) in the marmoset. On the medial wall, the putative marmoset network includes anterior cingulate areas 32, 25 and 24 and posterior cingulate areas 29a-c, 29d, 23a, 23b, PGM, the prostriate area (ProSt) and area TH. Subcortically, we observed the involvement of the caudate, different portions of the thalamus (ventral anterior, mediodorsal and lateral geniculate nuclei) and the superior colliculus.

Finally, we investigated the theory of mind (TOM) network, one of the highest-level cognitive networks in humans, who involves areas relevant for inferring others mental states(51–53). The mask we selected, largely overlapping with the more “cognitive” nuance of the TOM described by Schurz and colleagues(53), includes an occipito-temporal cluster (PG, TPOJ, STV, PFm), superior temporal sulcus regions (STS, TE1, TE2, TG, STGa), and a prefrontal cluster (areas 45, 47l, 8aD and 9a), with additional involvement of medial wall regions (prefrontal areas 32, 25, 24, 10 and 9 and posterior cingulate areas 23v, 31 and 7m) and subcortical regions (cerebellum, hippocampus, amygdala, caudate). This network is confirmed through within-species correlation, which revealed a pattern of areas predominantly limited to the previously described mask, with the addition of some lower correlations in the dorsal prefrontal cortex and, subcortically, in the anterior portions of the thalamus and of the caudate nucleus. The correlation of this network with the marmoset brain reveals only a sparse pattern of activations. Some correlation spots seem to reflect those observed in the human sample, such as the cluster at the occipito-temporo-parietal junction (MST, FST) or those located along the superior temporal sulcus and the temporal pole (TE3, TE1, areas 35 and 36). Similarly, the medial wall of the putative marmoset TOM network includes a more anterior cluster (areas 32 and 24) and a more posterior one (coinciding with area 29). Additional correlations are observed in the prefrontal/premotor cortex (areas 8aD, 13, and the dorsal area 6) and in the occipital cortex. Subcortically, the correlation map included the most anterior part of the caudate, the putamen and the lateral nucleus of the amygdala. These results are displayed in Figure 5C.

### Motor networks

In the motor domain, we searched for marmoset homologues of three human networks: the reaching/grasping, the saccade, and the speech production network. For the same reasons as described above, only the results concerning the right hemisphere are represented in this study for the reaching/grasping and saccades networks, while only the left hemisphere is represented for the speech production network (due to the well-known left lateralization of speech in humans(54,55)).

The human reaching/grasping network encompasses areas involved in the planning and performance of reaching and grasping movements(56). This network includes areas in the intraparietal sulcus (MIP, LIP, AIP, VIP), area 7PC, the PF complex, and the somatosensory, motor and premotor cortices. Moreover, two clusters of activations are observed on the median wall, one in the parieto-occipital junction (7Am, 7Pm, 7m, DVT, POS2 and V6) and one in the cingulate cortex, including areas 24, 23, 33 and the SCEF. The within-species correlation of this network is mainly consistent with the described pattern, although with lower correlations of the somato-motor areas and stronger correlations extending across the cingulate cortex. The putative marmoset reaching-grasping network shows correlation clusters in the superior visual areas and the intraparietal sulcus (V6, V6A, MIP, VIP, LIP, OPt, PEC), the prefrontal/premotor cortex (6DR and area 8), and multiple regions of the cingulate cortex (32, 32v, and 9 in the anterior part, 30 and 29 in the central and posterior parts). In addition to these clusters, which align with the human mask, there are two extensive clusters in the occipito-temporal region (see Figure 6A). Subcortically, a robust correlation spot emerges in the amygdala.

**Figure 6.**
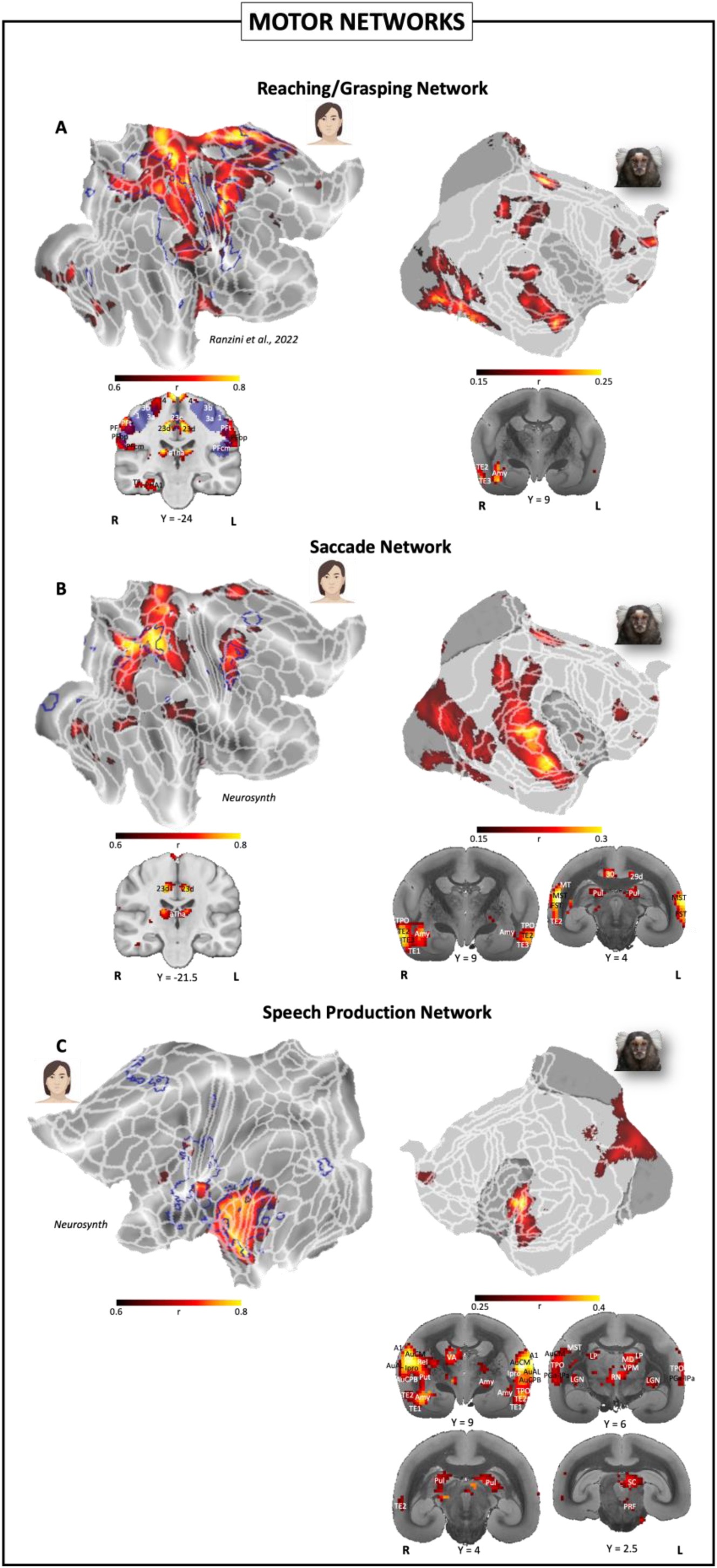
Motor functional network correlation between human and marmosets. The figure displays the human-marmoset correlations concerning A) the reaching/grasping network, B) the saccades network, and C) the speech production network. In the left column of each panel, the masks used to extract the network timecourse from our human sample (blue colour) are displayed on flat maps of the right hemisphere. Hot colour represents the within-species correlation of the timecourse of the network with all the voxels of the brain. White lines delineate the multi-modal cortical parcellation atlas(36). In the right column, the correlation maps obtained correlating the human network timecourses to the marmoset brain timecourse, on a voxel-by-voxel basis, are displayed on flat maps of the right hemisphere. White lines delineate the Paxinos parcellation(37) of the NIH marmoset brain atlas(34). For all maps, the volumetric cortical and subcortical activations at different inter-aural levels are displayed on the coronal slices of MR anatomical images of a human (left column) or a marmoset (right column) brain.

In humans, the network underlying saccade generation includes the intraparietal sulcus (VIP, MIP, LIP), visual areas V1, V3A-B and V7, the primary motor cortex, and a strong precentral cluster involving FEF, PEF, 55b, 6d, 6a and 6v. On the median wall, a cluster of activation includes the SCEF. The within-species correlation confirmed this pattern of areas and revealed additional strong correlations in the posterior parietal cortex (areas 7, 5, and precuneus) and the cingulate cortex (particularly area 23). Subcortically, the involvement of the anterior portion of the thalamus is observed. The between-species correlation showed a frontal correlation cluster (6DR, 8c, 8aV, and 8aD - where the marmoset FEF is located(57)), a cluster in the posterior cingulate cortex (areas 29 and 30), and an extensive correlation cluster extending from the occipito-parietal region (V6, V6A, MIP, LIP, VIP, OPt, 19DI) to the temporal region (MT, V4t, MST, FST, PGa-IPa, TPO, TE3, TE2, TE1). Subcortically, the recruitment of the amygdala and the pulvinar is observed bilaterally. These results are depicted in Figure 6B.

Lastly, we looked for a functional homologue of the human speech production network. In humans, this network includes auditory cortex and the superior temporal sulcus, as in the vocalization processing network. In addition, it includes somatosensory and motor cortices, premotor regions, and cingulate areas (24dv, 24pr, 32pr and the supplementary and cingulate eye field, SCEF). Subcortically, the network included portions of the cerebellum, the amygdala, and the thalamus. Given the absence of a language production component in our study, it is not surprising that the within-species correlation revealed a pattern of areas primarily confined to the auditory and insular regions, with only a few limited correlation regions in the somatosensory areas. The subcortical structures present in the initial mask are also absent. In the same way, the correlation of this network timecourse with the marmoset brain timecourse revealed a network encompassing auditory (core, belt and parabelt areas), insular, temporal (TPO, STR, and the conjunction between TE1, TE2, TE3 and 36) and occipital areas, as showed in Figure 6C. On the medial wall we observed correlations in area 32, while subcortically the amygdala, SC, pulvinar and several thalamic nuclei were correlated. No reliable correlation was observed in somatosensory, motor, or premotor cortices.

### Limbic network

The final network that we explored was the human fear network, considered as the system of areas activated during the feeling of fear(58–60). As for the previous networks, only the results concerning the right hemisphere are described in this manuscript.

The human fear network includes both cortical areas and limbic structures. As shown in Figure 7, these regions include the ventral portion of the temporal lobe (FFC, entorhinal cortex, PeEc, STGa), some areas on the medial wall (25, 24, 32 and 10) and subcortical structures (amygdala, hippocampus and posterior thalamus). The within-species correlation, despite the absence of frightening elements in the experimental material, revealed a pattern of areas consistent with the described findings, showing correlations with key structures of the fear network, such as the amygdala. Lower correlations are also observed in the STS, the frontal cortex (FEF, 55b, PEF, inferior frontal gyrus), and the dorsomedial prefrontal cortex (SCEF, 8BM, 32pr). When correlating the timecourse of this network with the marmoset voxel timecourses, we obtained a correlation map including clusters in the visual (V2, V3), temporal (TE1, 35, 36) and parietal cortices (LIP, VIP, OPt), several areas of the medial wall of the brain (23, 30, PG and 32) and subcortical structures such as the lateral geniculate nucleus, the amygdala, the caudate and the putamen. These results are illustrated in Figure 7.

**Figure 7.**
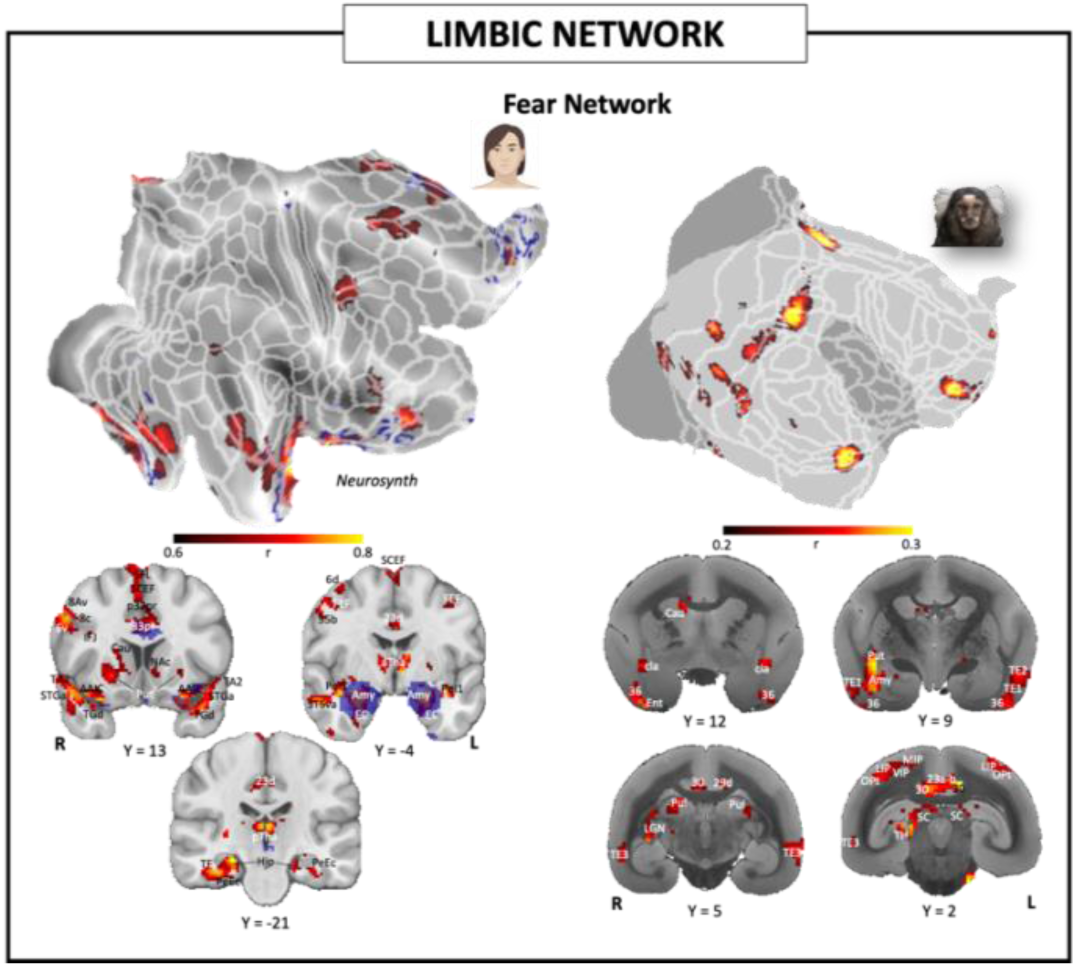
Limbic functional network correlation between human and marmosets. The figure displays the human-marmoset correlations concerning the fear network. In the left column, the mask used to extract the network timecourse from our human sample (blue colour) is displayed on flat map of the right hemisphere. Hot colour represents the within-species correlation of the timecourse of the network with all the voxels of the brain. White lines delineate the multi-modal cortical parcellation atlas(36). In the right column, the correlation map obtained correlating the human network timecourses to the marmoset brain timecourse, on a voxel-by-voxel basis, is displayed on flat map of the right hemisphere. White lines delineate the Paxinos parcellation(37) of the NIH marmoset brain atlas(34). For all maps, the volumetric cortical and subcortical activations at different inter-aural levels are displayed on the coronal slices of MR anatomical images of a human (left column) or a marmoset (right column) brain.

### Coincidence map

In the final stage of our analysis, we computed a coincidence map to identify brain regions in humans and marmosets that consistently correlate with the functional networks we examined (Figure 8). For clarity, this study presents unilateral results; however, the bilateral coincidence map offers a comprehensive view of correlation distributions across both hemispheres.

**Figure 8.**
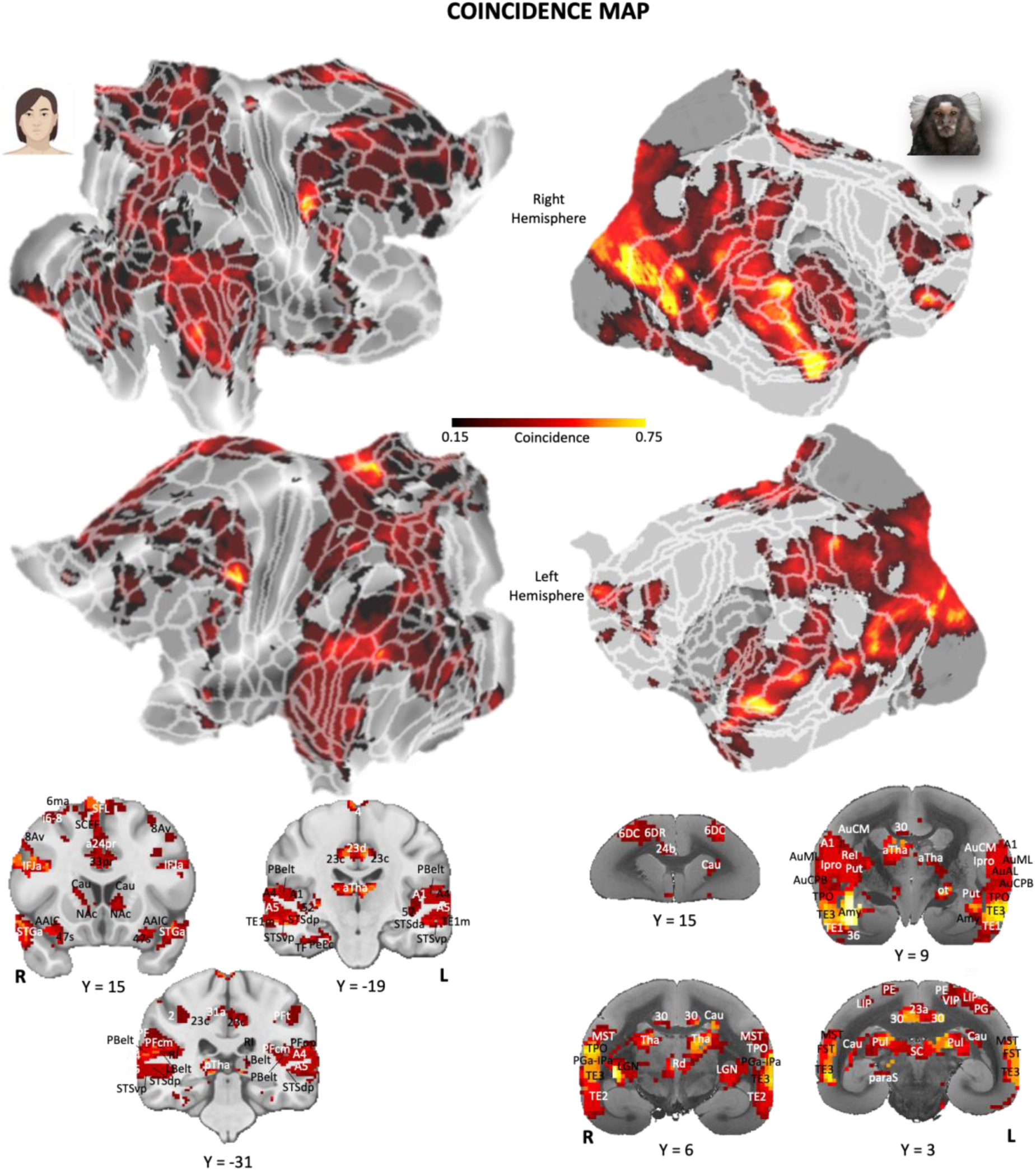
Coincidence of the correlation maps in human and marmosets. The figure displays the overlap of the within-species (human-to-human) and between-species (human-to-marmoset) correlation maps. Coincidence values range from 0 (voxels never included in the correlation maps) to 1 (voxels over-threshold in all correlation maps). In the left column, the coincidence of the human within-species correlation maps is displayed on flat maps of both hemispheres. White lines delineate the multi-modal cortical parcellation atlas(36). In the right column, the same variable is displayed on flat maps for the marmoset between-species correlation maps. For both maps, the volumetric cortical and subcortical activations at different inter-aural levels are displayed on the coronal slices of MR anatomical images of a human (left column) or a marmoset (right column) brain.

In our human subjects (Figure 8, left part), prominent coincidence peaks were observed in the precentral gyrus, encompassing the frontal eye field (FEF), area 55b, and the parieto-eye field (PEF), bilaterally. Another significant region of high coincidence was the posterior cingulate cortex, which includes areas 23c, 5mv, and the precuneus. Additionally, the functional networks predominantly engaged brain regions along the temporo-parieto-occipital junction and the superior temporal sulcus (STS). Subcortically, the anterior thalamus was the most recurrently recruited structure. These clusters suggest the presence of multifunctional hubs, reflecting the broad brain coverage achieved by our experimental methodology, which includes occipital, temporal, parietal, frontal, and prefrontal regions. Less prominent in our findings were the somatosensory, motor, olfactory, gustatory, and orbitofrontal regions.

For the marmoset subjects, as illustrated in Figure 8 (right part), the coincidence mapping shows greater and more extensive overlap than in the human sample. The most pronounced areas of overlap were along the occipito-temporal axis, including both primary visual areas and areas MST, FST, PGa-IPa, TPO, and the junction of areas TE1, 35, and 36. Additional significant coincidence peaks were observed at the junction between prefrontal areas 13 and 11, and in the posterior and anterior cingulate cortex (areas 29, 30, and 32 respectively). Subcortical structures such as the superior colliculus, amygdala, and various thalamic nuclei also showed frequent involvement in the correlation maps. Similar to the human findings, the marmoset results underscore the extensive recruitment of cerebral cortical and subcortical structures by the selected networks, albeit with under-recruitment of somato-motor, olfactory, and gustatory areas.

## Discussion

We employed movie-driven functional magnetic resonance imaging (md-fMRI) to capture BOLD responses in both humans and marmosets exposed to a naturalistic movie featuring diverse visual (marmosets, humans, macaques, natural landscapes, urban environments) and auditory stimuli (human speech, marmoset calls, other animal vocalizations, and music). Building on our prior use of md-fMRI to investigate functional homologies between marmosets and humans(28), particularly in high-order visual areas involved in face processing, this study broadens our inquiry by integrating auditory stimuli, increasing the range of cortical and subcortical brain areas activated in both species. The enhanced stimulation design allows us to identify networks that are potential functional homologues to well-characterized long-range human networks, thereby providing deeper insights into the neural architecture and functional capacities of marmosets relative to humans.

A key finding is the robust correlation between the activity in the human MT area (hMT) and its marmoset counterpart (mMT), known for processing visual motion in both species(61–64). The consistent MT activity correlation across species underscored the validity of our experimental paradigm. Additionally, the functional connectivity observed between mMT and adjacent areas (FST, MST, V4t)(65), aligns with the correlated activity between hMT and the broader marmoset brain (Figure 1), lending further credence to our interpretations.

Interestingly, the human-to-marmoset correlation map extends significantly across the marmoset temporal lobe—a pattern not mirrored in the human-to-human correlation map. However, in our human subjects, the connectivity of hMT broadly overlaps with regions activated by networks involved in face processing(16,18,21) and action observation(20,66). Similarly, extensive occipito-temporal clusters appear when correlating the activity of these human networks (but also other networks with significant visual components, as the places and audiovisual networks) with the marmoset brain timecourse. This suggests a lower specialization in marmoset occipito-temporal areas compared to humans, potentially implying that different neural populations might be involved in processing diverse functions—such as face recognition, action observation, environmental scanning, and audiovisual integration— although such fine segregation may not be discernible with fMRI technology. Alternatively, this cluster’s strong association with the processing of complex, dynamic visual stimuli, like faces and actions in the movie, could indicate a high degree of interconnectedness with the MT region and its periphery.

Exploring the Action Observation Network (AON) in more depth reveals how md-fMRI aligns consistently with existing human(39–41) and marmoset(20) fMRI studies. Notably, the occipito-temporal and prefrontal nodes of the AON are present in the literature of both species, whereas the inferior parietal lobule, typically part of the human AON, does not feature in the marmoset network. This pattern of results is fully replicated here by correlating the human AON timecourse with the marmoset one, supporting our interpretations. Intriguingly, the prefrontal cluster is less pronounced in md-fMRI compared to traditional localizer paradigms. This discrepancy may stem from the nature of the stimuli used: whereas localizer paradigms specifically target the AON with direct stimuli, a naturalistic movie introduces a broader array of stimuli, which often occur simultaneously, potentially diluting the localized activation typically seen in more specific experimental setups.

Our findings on the face processing network corroborate existing marmoset fMRI research(16–18,21,28), revealing an extensive occipito-temporal cluster including previously identified marmoset face-patches(16,18), and a localized prefrontal cluster. This observation aligns with our group’s recent discoveries of a marmoset face-patch at the intersection of area 8Av and area 45(16,21). Moreover, the potential functional homology between the human perirhinal cortex (PeEc) and marmoset area 36, as previously speculated(28), is further substantiated in this study.

Concerning the places network, the distinct patch at the convergence of areas V4, TEO, and TFO may represent the marmoset equivalent of the "lateral place patch" identified in macaques(67). Additionally, a significant patch in the posterior cingulate cortex appears to correspond to the location of the retrosplenial complex (or medial place area), known for its specificity to place processing in both humans and macaques(45,68). Notably, this patch is absent from the other two human visual networks we analyzed and is not part of the corresponding marmoset networks, indicating its unique role in scene processing.

The three visual networks analyzed share a common correlation cluster in prefrontal area 8aV, which is the location of the marmoset FEF(57). The FEF is implicated not only in saccade execution but also in directing both overt and covert spatial attention(69). Our findings reaffirm the localization and functional role of the FEF in marmosets; this area appears consistently in correlation maps of networks that necessitate saccade execution and the orientation of visual spatial attention. Notably, the FEF is present in the correlation maps for the saccades network, the reaching/grasping network, and the central executive network. Conversely, it is absent from the correlation maps of networks not involving attentional processes, such as the audiovisual integration network. In this last multisensory network, pronounced correlation clusters are evident along the occipito-temporal axis and within the auditory cortex, confirming the previous marmoset literature(21). A prefrontal cluster is also present, situated at the junction between areas 13M, 13L, and 11, a region reporting anatomical(70) and functional(71) connections with areas pivotal in multisensory integration. This patch is coherent with the cluster described by Dureux and colleagues, even if this latter extends further toward areas 47 and 8aV, probably due to the presentation of marmoset faces and vocalizations combined(21).

The auditory networks (vocalization, language comprehension and speech production, given the absence of a somato-motor component in our study) show strong functional homology in the auditory cortex, as expected. For the language comprehension, a purely human network, this correlation spot is almost the only functional homology observable. Surprisingly, the motor network for speech production shows more extensive correlations in the visual cortex, resembling the vocalization network. This is probably due to the visual patch of the speech production network mask we used, located between V4 and LO2. These correlations are thus explained by purely perceptual processes, and the lack of correlations at the somatosensory and motor levels instead confirms (and limits) our experimental method.

In contrast, the correlation of the vocalization network demonstrates excellent overlap with both the human fMRI literature(47,72,73) and our group’s previous results in marmosets(6,21). Only exception is the absence of somato-motor activations, described with our previous fMRI localizer(6) in marmosets and probably due to vocalization production, in an experimental paradigm crowded with conspecific vocalizations. This interpretation aligns with the pronounced vocal nature of common marmoset communication and their cooperative antiphonal calling(74). Here, where marmoset vocalizations were less frequent and interleaved with other auditory stimuli, it is likely that the perception of conspecific calls did not lead to antiphonal calls.

The confirmation of the involvement of mPFC area 32 in the marmoset vocalization network(6,21), and its presence in all the cognitive networks we investigated, support the hypothesis of this region being a crucial functional hub in the marmoset brain. The presence of area 32 in the marmoset’s default mode network (DMN) also addresses the previously missing mPFC patch in the marmoset DMN(75), a main component of the human(49,76), macaque(77–79), and even rodent DMN(80–82). Thus, area 32 may be a crucial hub for high-level cognitive processes in marmosets, warranting further studies to understand the functions of different neural populations in this region.

The parallel between species further includes the presence, in the marmoset, of a rostral and a posterior cingulate cluster, mirroring what observed in all human cognitive networks. While the exact human cluster locations differ slightly, the correlated marmoset areas overlap extensively, suggesting reduced cortical specialization in marmosets. This is also evident in the prefrontal cortex, where human cognitive networks show activation clusters correlating with overlapping marmoset prefrontal areas. Interestingly, the temporal and occipito-temporal activation clusters of the human TOM network found potentially homologues along the marmoset STS and at the junction between MST and FST areas. It is surprising how a human cognitive network involved in ascribing mental states to others(51,52) maps to the marmoset, given that the fMRI protocol materials were not designed to trigger theory of mind processes. However, these brain regions likely participate in various processes and may be similarly recruited during movie viewing in both species. Combined with findings that marmosets and humans activate comparable brain networks(8) when viewing Frith-Happé animations(83,84), however, this suggests the potential presence of a TOM network in the common marmoset.

Concerning the human network underlying executive functions, it is necessary to specify that multiple labels are used to denote slightly different patterns of fronto-parietal brain areas(50). In addition to the functional parallels in the anterior (ACC) and posterior cingulate cortex (PCC), the human central executive network we selected(50) reported broad correlation in the prefrontal cortex of the marmoset. This is a remarkable example of the validity of this movie-driven fMRI approach in establishing functional homologies across species. Recently, single neuron recordings have reported delayed-related activity in prefrontal areas in marmosets during a delay-match-to-position task(85), consistent with the well-known role of the prefrontal cortex in working memory in humans(50) and macaque monkeys(86) and further supporting our present findings.

An interesting aspect observed through the correlation of cognitive networks is the pervasive presence of a correlation cluster in the marmoset’s posterior parietal cortex, roughly corresponding to the areas of the intraparietal sulcus VIP, MIP, and LIP, plus the area Opt. This activation cluster is not present in the human networks we investigated but is noted in marmoset fMRI literature concerning the DMN(11,75), TOM(8) and attention network(11). This cluster likely correlates with multiple human cognitive networks, as the posterior parietal region in marmosets serves as a functional hub involved in various roles, both during external information processing, multisensory integration and rest(9–11,21,75,87,88). The interpretation of this cluster’s function becomes more complex considering that the marmoset’s posterior parietal cortex is involved in saccade-related activity, as demonstrated through fMRI(7), single-unit recordings(89), and electrical microstimulation(90). This is confirmed by our results, as the correlation of the human saccade network with the marmoset timecourse reveals a cluster in the posterior parietal cortex centered on the LIP area.

The functional homologies between this human network and the marmoset saccade network continues with the correlation of the prefrontal area 8aV, where the marmoset FEF is located(57). The human saccade network appears to be preserved in the marmoset, although a broad correlation cluster along the occipito-temporal axis is observed. This is likely due to the presence of activation patches in primary (V1, V2) and higher-level visual areas (V3, V7) within the human network we considered.

Regarding a second motor network considered, the reaching/grasping network(56), the results obtained through correlation reveal both a limitation and a confirmation of the md-fMRI methodology. The lack of complete correlation of this network is indeed predictable, given the lack of motor activity within the scanner in our experiment. On the other hand, this lack underscores the impossibility of investigating the functional homologies between the two species at the level of motor and somatosensory areas using md-fMRI. Such a correlation, however, is not impossible but requires a more specific task-based experimental paradigm and a training path for non-human primates.

Finally, the correlation of the timecourse of the only limbic network we investigated highlights the effectiveness of our experimental paradigm in identifying functional homologies even at subcortical levels. Notably, the absence of specific stimuli capable of activating the fear network concurrently in humans and marmosets may pose a challenge to the computation of robust cross-species correlations. Taking this into account, it is surprising to observe the functional homologies between the human fear network and its correlated counterpart in the marmoset. This includes the amygdala, a prominent subcortical component of the human fear network(59,91,92), bilaterally. Additionally, the human activations in PeEc and the entorhinal cortex seems to find correspondence in the putative homologous area 36 of the marmoset(28). On the medial wall, the correlation again highlights the recruitment of area 32, possibly corresponding to the human fear network patch in the ACC, and a correlation in the PCC, potentially corresponding to the human activation cluster located between areas 24pr and 33pr. Finally, a correlation cluster is again observed in the posterior parietal cortex of the marmoset, overlapping with those already described for cognitive networks and not included in the human fear network.

The coincidence map provides a compelling overview of our findings. Notably, both the human within-species correlation maps and the marmoset between-species maps show two peaks at the posterior cingulate cortex and the FEF. This overlap is likely due to our selection of networks, which include multiple visual and cognitive networks that recruit these regions. However, the functional parallelism observed between the two species is surprising, considering that no specific network was targeted in our experimental material.

Another common point between humans and marmosets is the extensive recruitment of the occipito-temporal region. This not only demonstrates functional parallelism but also confirms previous marmoset fMRI literature. Ghahremani and colleagues have shown that the occipito-temporal axis, along with the dorsal prefrontal cortex and the orbitofrontal area, is one of the most important functional hubs in the common marmoset(10). These key points of the marmoset brain’s functional architecture are involved in many of the functional networks we selected, as evidenced by the coincidence map.

In conclusion, the use of md-fMRI with the same naturalistic movie presented to two different primate species has allowed us to identify putative functional homologies of well-known human functional networks in the common marmoset. These findings show a high degree of consistency with both the fMRI literature on humans and previous task-based or resting-state fMRI results on marmosets, where available. Additionally, we have made all the human and marmoset data and code fully available. The main objective of this study is to provide other research groups the opportunity to test functional networks beyond those we investigated and to extend the cross-species comparison between humans and the common marmoset.

### Limitations

While our current md-fMRI approach may be too varied for precise functional localization studies, it is possible to create more selective movies focusing on specific features, such as multiple and more frequent conspecific calls or varying levels of motion or contrast. Additionally, targeting specific brain regions or functional patches could further investigate functional homologies between species or analyze responses during specific periods of the movie, such as correlating the human DMN only during baseline periods or the AON during action scenes.

The absence of motor, somatosensory, olfactory, and gustatory areas in the coincidence map was an anticipated limitation of our experimental method, as it lacks elements or stimuli capable of reliably activating these areas.

Moreover, a significant limitation of our approach is that brain regions with closely interconnected functions, such as those along the marmoset’s occipito-temporal axis, are likely to be clustered together due to extensive timecourse overlap. In cases where more in-depth and localized analysis is required to distinguish various functional populations, alternative techniques such as functional localizers or electrophysiological recordings, or the use of more specific movies, must be employed.

## Acknowledgements

Support was provided by a Discovery grant by the Natural Sciences and Engineering Research Council of Canada and the Canadian Institutes of Health Research (FRN 183973). We also acknowledge the support of the Government of Canada’s New Frontiers in Research Fund (NFRF), [NFRF-T-2022-00051]. We wish to thank Cheryl Vander Tuin, Whitney Froese, Hannah Pettypiece, and Miranda Bellyou for animal preparation and care, Dr. Alex Li and Trevor Szekeres for scanning assistance, Dr. Kyle Gilbert, and Peter Zeman for coil designs.

## Data Availability

The group and individual maps for the human and marmoset samples, along with the code underpinning this study, are fully available online at 10.5281/zenodo.12746414.

## Supplementary Information

**Supplementary Figure 1.**
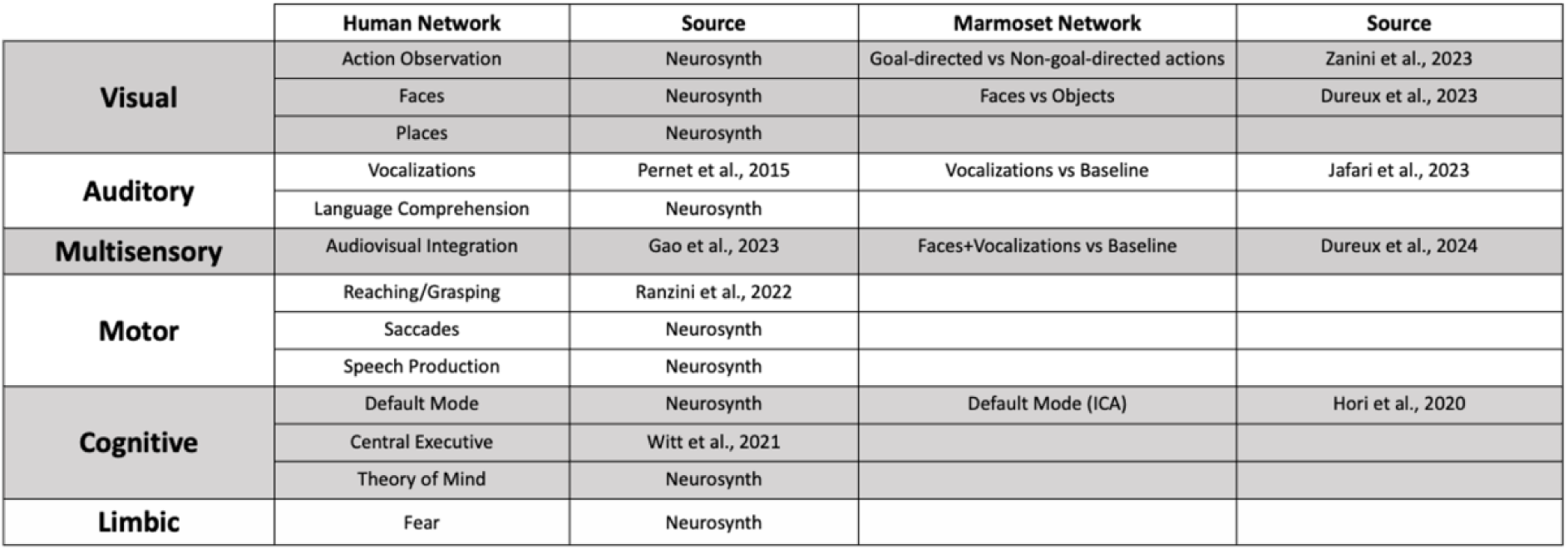
Functional large-scale networks investigated in this study. Each row represents a distinct functional network. In the first two columns, the human functional network and the source of the mask utilized in this study, respectively. In the last two columns, when available, the marmoset functional map used as a comparison and the relative source.

## References

1. Okano H. Current Status of and Perspectives on the Application of Marmosets in Neurobiology. Annual Review of Neuroscience. 2021;44(1):27–48.

2. Burman KJ, Rosa MGP. Architectural subdivisions of medial and orbital frontal cortices in the marmoset monkey (Callithrix jacchus). Journal of Comparative Neurology. 2009;514(1):11–29.

3. Solomon SG, Rosa MGP. A simpler primate brain: the visual system of the marmoset monkey. Front Neural Circuits [Internet]. 2014 Aug 8 [cited 2024 Apr 9];8. Available from: https://www.frontiersin.org/articles/10.3389/fncir.2014.00096

4. Chen CY, Matrov D, Veale R, Onoe H, Yoshida M, Miura K, et al. Properties of visually guided saccadic behavior and bottom-up attention in marmoset, macaque, and human. Journal of Neurophysiology. 2021 Feb;125(2):437–57.

5. Mitchell JF, Leopold DA. The marmoset monkey as a model for visual neuroscience. Neuroscience Research. 2015 Apr 1;93:20–46.

6. Jafari A, Dureux A, Zanini A, Menon RS, Gilbert KM, Everling S. A vocalization-processing network in marmosets. Cell Reports. 2023 May 30;42(5):112526.

7. Schaeffer DJ, Gilbert KM, Hori Y, Hayrynen LK, Johnston KD, Gati JS, et al. Task-based fMRI of a free-viewing visuo-saccadic network in the marmoset monkey. NeuroImage. 2019 Nov 15;202:116147.

8. Dureux A, Zanini A, Selvanayagam J, Menon RS, Everling S. Gaze patterns and brain activations in humans and marmosets in the Frith-Happé theory-of-mind animation task. Elife. 2023 Jul 14;12:e86327.

9. Garin CM, Hori Y, Everling S, Whitlow CT, Calabro FJ, Luna B, et al. An evolutionary gap in primate default mode network organization. Cell Reports. 2022 Apr 12;39(2):110669.

10. Ghahremani M, Hutchison RM, Menon RS, Everling S. Frontoparietal Functional Connectivity in the Common Marmoset. Cerebral Cortex. 2017 Aug 1;27(8):3890–905.

11. Hori Y, Schaeffer DJ, Yoshida A, Cléry JC, Hayrynen LK, Gati JS, et al. Cortico-Subcortical Functional Connectivity Profiles of Resting-State Networks in Marmosets and Humans. J Neurosci. 2020 Nov 25;40(48):9236–49.

12. Ji JL, Spronk M, Kulkarni K, Repovš G, Anticevic A, Cole MW. Mapping the human brain’s cortical-subcortical functional network organization. NeuroImage. 2019 Jan 15;185:35–57.

13. Yeo TBT, Krienen FM, Sepulcre J, Sabuncu MR, Lashkari D, Hollinshead M, et al. The organization of the human cerebral cortex estimated by intrinsic functional connectivity. Journal of Neurophysiology. 2011 Sep;106(3):1125–65.

14. Pinsk MA, Arcaro M, Weiner KS, Kalkus JF, Inati SJ, Gross CG, et al. Neural Representations of Faces and Body Parts in Macaque and Human Cortex: A Comparative fMRI Study. Journal of Neurophysiology. 2009 May;101(5):2581–600.

15. Laurent MA, Audurier P, De Castro V, Gao X, Durand JB, Jonas J, et al. Towards an optimization of functional localizers in non-human primate neuroimaging with (fMRI) frequency-tagging. NeuroImage. 2023 Apr 15;270:119959.

16. Dureux A, Zanini A, Everling S. Face-Selective Patches in Marmosets Are Involved in Dynamic and Static Facial Expression Processing. J Neurosci. 2023 May 10;43(19):3477–94.

17. Schaeffer DJ, Selvanayagam J, Johnston KD, Menon RS, Freiwald WA, Everling S. Face selective patches in marmoset frontal cortex. Nat Commun. 2020 Sep 25;11(1):4856.

18. Hung CC, Yen CC, Ciuchta JL, Papoti D, Bock NA, Leopold DA, et al. Functional Mapping of Face-Selective Regions in the Extrastriate Visual Cortex of the Marmoset. J Neurosci. 2015 Jan 21;35(3):1160–72.

19. Hung CC, Yen CC, Ciuchta JL, Papoti D, Bock NA, Leopold DA, et al. Functional MRI of visual responses in the awake, behaving marmoset. NeuroImage. 2015 Oct 15;120:1–11.

20. Zanini A, Dureux A, Selvanayagam J, Everling S. Ultra-high field fMRI identifies an action-observation network in the common marmoset. Commun Biol. 2023 May 22;6(1):1–11.

21. Dureux A, Zanini A, Everling S. Mapping of facial and vocal processing in common marmosets with ultra-high field fMRI. Commun Biol. 2024 Mar 13;7(1):1–15.

22. Hasson U, Furman O, Clark D, Dudai Y, Davachi L. Enhanced Intersubject Correlations during Movie Viewing Correlate with Successful Episodic Encoding. Neuron. 2008 Feb 7;57(3):452–62.

23. Hasson U, Malach R, Heeger DJ. Reliability of cortical activity during natural stimulation. Trends in Cognitive Sciences. 2010 Jan 1;14(1):40–8.

24. Lerner Y, Honey CJ, Silbert LJ, Hasson U. Topographic Mapping of a Hierarchy of Temporal Receptive Windows Using a Narrated Story. J Neurosci. 2011 Feb 23;31(8):2906–15.

25. Naci L, Cusack R, Anello M, Owen AM. A common neural code for similar conscious experiences in different individuals. Proceedings of the National Academy of Sciences. 2014 Sep 30;111(39):14277–82.

26. Russ BE, Leopold DA. Functional MRI mapping of dynamic visual features during natural viewing in the macaque. NeuroImage. 2015 Apr 1;109:84–94.

27. Park SH, Koyano KW, Russ BE, Waidmann EN, McMahon DBT, Leopold DA. Parallel functional subnetworks embedded in the macaque face patch system. Science Advances. 2022 Mar 9;8(10):eabm2054.

28. Hori Y, Cléry JC, Selvanayagam J, Schaeffer DJ, Johnston KD, Menon RS, et al. Interspecies activation correlations reveal functional correspondences between marmoset and human brain areas. Proceedings of the National Academy of Sciences. 2021 Sep 14;118(37):e2110980118.

29. Gilbert KM, Dureux A, Jafari A, Zanini A, Zeman P, Menon RS, et al. A radiofrequency coil to facilitate task-based fMRI of awake marmosets. J Neurosci Methods. 2023 Jan 1;383:109737.

30. Zanini A, Dureux A, Jafari A, Gilbert KM, Zeman P, Bellyou M, et al. *In vivo* functional brain mapping using ultra-high-field fMRI in awake common marmosets. STAR Protocols. 2023 Dec 15;4(4):102586.

31. Gilbert KM, Klassen LM, Mashkovtsev A, Zeman P, Menon RS, Gati JS. Radiofrequency coil for routine ultra-high-field imaging with an unobstructed visual field. NMR in Biomedicine. 2021 Mar 1;34(3).

32. Cox RW. AFNI: Software for Analysis and Visualization of Functional Magnetic Resonance Neuroimages. Computers and Biomedical Research. 1996 Jun;29(3):162–73.

33. Smith SM, Jenkinson M, Woolrich MW, Beckmann CF, Behrens TEJ, Johansen-Berg H, et al. Advances in functional and structural MR image analysis and implementation as FSL. NeuroImage. 2004 Jan 1;23:S208–19.

34. Liu C, Ye FQ, Yen CCC, Newman JD, Glen D, Leopold DA, et al. A digital 3D atlas of the marmoset brain based on multi-modal MRI. NeuroImage. 2018 Apr 1;169:106–16.

35. Avants BB, Tustison NJ, Song G. Advanced Normalization Tools (ANTs). Insight J. 2009;2(365):1–35.

36. Glasser MF, Coalson TS, Robinson EC, Hacker CD, Harwell J, Yacoub E, et al. A multi-modal parcellation of human cerebral cortex. Nature. 2016 Aug;536(7615):171–8.

37. Paxinos G, Watson C, Petrides M, Rosa MGP, Tokuno H. The marmoset brain in stereotaxic coordinates. Elsevier Academic Press; 2012.

38. Saleem KS, Avram AV, Glen D, Schram V, Basser PJ. The Subcortical Atlas of the Marmoset (“SAM”) monkey based on high-resolution MRI and histology [Internet]. bioRxiv; 2024 [cited 2024 Mar 27]. p. 2024.01.06.574429. Available from: https://www.biorxiv.org/content/10.1101/2024.01.06.574429v1

39. Caspers S, Zilles K, Laird AR, Eickhoff SB. ALE meta-analysis of action observation and imitation in the human brain. NeuroImage. 2010 Apr 15;50(3):1148–67.

40. Cross ES, Hamilton AF de C, Kraemer DJM, Kelley WM, Grafton ST. Dissociable substrates for body motion and physical experience in the human action observation network. European Journal of Neuroscience. 2009;30(7):1383–92.

41. Gazzola V, Rizzolatti G, Wicker B, Keysers C. The anthropomorphic brain: The mirror neuron system responds to human and robotic actions. NeuroImage. 2007 May 1;35(4):1674–84.

42. Kanwisher N, McDermott J, Chun MM. The Fusiform Face Area: A Module in Human Extrastriate Cortex Specialized for Face Perception. J Neurosci. 1997 Jun 1;17(11):4302–11.

43. Haxby JV, Hoffman EA, Gobbini MI. The distributed human neural system for face perception. Trends in Cognitive Sciences. 2000 Jun 1;4(6):223–33.

44. Posamentier MT, Abdi H. Processing Faces and Facial Expressions. Neuropsychol Rev. 2003 Sep 1;13(3):113–43.

45. Epstein RA, Baker CI. Scene perception in the human brain. Annu Rev Vis Sci. 2019 Sep 15;5:373–97.

46. Henderson JM, Hollingworth A. High-level scene perception. Annual Review of Psychology. 1999 Feb 1;50(Volume 50, 1999):243–71.

47. Pernet CR, McAleer P, Latinus M, Gorgolewski KJ, Charest I, Bestelmeyer PEG, et al. The human voice areas: Spatial organization and inter-individual variability in temporal and extra-temporal cortices. NeuroImage. 2015 Oct 1;119:164–74.

48. Gao C, Green JJ, Yang X, Oh S, Kim J, Shinkareva SV. Audiovisual integration in the human brain: a coordinate-based meta-analysis. Cerebral Cortex. 2023 May 1;33(9):5574–84.

49. Menon V. 20 years of the default mode network: A review and synthesis. Neuron. 2023 Aug 16;111(16):2469–87.

50. Witt ST, van Ettinger-Veenstra H, Salo T, Riedel MC, Laird AR. What Executive Function Network is that? An Image-Based Meta-Analysis of Network Labels. Brain Topogr. 2021 Sep 1;34(5):598–607.

51. Carruthers P, Smith PK. Theories of Theories of Mind. Cambridge University Press; 1996. 422 p.

52. Premack D, Woodruff G. Does the chimpanzee have a theory of mind? Behavioral and Brain Sciences. 1978 Dec;1(4):515–26.

53. Schurz M, Radua J, Tholen MG, Maliske L, Margulies DS, Mars RB, et al. Toward a hierarchical model of social cognition: a neuroimaging meta-analysis and integrative review of empathy and theory of mind. Psychological Bulletin [Internet]. 2020 [cited 2024 Jul 25];147(3). Available from: https://ora.ox.ac.uk/objects/uuid:00f31bea-f706-4182-ac8e-e9389cdf664b

54. Narain C, Scott SK, Wise RJS, Rosen S, Leff A, Iversen SD, et al. Defining a Left-lateralized Response Specific to Intelligible Speech Using fMRI. Cerebral Cortex. 2003 Dec 1;13(12):1362–8.

55. Springer JA, Binder JR, Hammeke TA, Swanson SJ, Frost JA, Bellgowan PSF, et al. Language dominance in neurologically normal and epilepsy subjects: A functional MRI study. Brain. 1999 Nov 1;122(11):2033–46.

56. Ranzini M, Scarpazza C, Radua J, Cutini S, Semenza C, Zorzi M. A common neural substrate for number comparison, hand reaching and grasping: A SDM-PSI meta-analysis of neuroimaging studies. Cortex. 2022 Mar 1;148:31–67.

57. Selvanayagam J, Johnston KD, Schaeffer DJ, Hayrynen LK, Everling S. Functional Localization of the Frontal Eye Fields in the Common Marmoset Using Microstimulation. J Neurosci. 2019 Nov 13;39(46):9197–206.

58. Kirstein C, Güntürkün O, Ocklenburg S. Ultra-high field imaging of the amygdala - A narrative review. Neuroscience and biobehavioral reviews [Internet]. 2023 Sep [cited 2024 Jun 19];152. Available from: https://pubmed.ncbi.nlm.nih.gov/37230235/

59. LeDoux JE. Emotion Circuits in the Brain. Annual Review of Neuroscience. 2000 Mar 1;23(Volume 23, 2000):155–84.

60. LeDoux JE. Coming to terms with fear. Proceedings of the National Academy of Sciences. 2014 Feb 25;111(8):2871–8.

61. Born RT, Bradley DC. STRUCTURE AND FUNCTION OF VISUAL AREA MT. Annual Review of Neuroscience. 2005 Jul 21;28(Volume 28, 2005):157–89.

62. DeYoe EA, Essen DCV. Concurrent processing streams in monkey visual cortex. Trends in Neurosciences. 1988;11(5):219–26.

63. Lui LL, Rosa MGP. Structure and function of the middle temporal visual area (MT) in the marmoset: Comparisons with the macaque monkey. Neuroscience Research. 2015 Apr 1;93:62–71.

64. Saito H, Yukie M, Tanaka K, Hikosaka K, Fukada Y, Iwai E. Integration of direction signals of image motion in the superior temporal sulcus of the macaque monkey. J Neurosci. 1986 Jan 1;6(1):145–57.

65. Abe H, Tani T, Mashiko H, Kitamura N, Hayami T, Watanabe S, et al. Axonal Projections From the Middle Temporal Area in the Common Marmoset. Front Neuroanat. 2018;12:89.

66. Suzuki W, Banno T, Miyakawa N, Abe H, Goda N, Ichinohe N. Mirror Neurons in a New World Monkey, Common Marmoset. Frontiers in Neuroscience [Internet]. 2015 [cited 2022 Jun 6];9. Available from: https://www.frontiersin.org/article/10.3389/fnins.2015.00459

67. Kornblith S, Cheng X, Ohayon S, Tsao DY. A network for scene processing in the macaque temporal lobe. Neuron. 2013 Aug 21;79(4):766–81.

68. Nasr S, Liu N, Devaney KJ, Yue X, Rajimehr R, Ungerleider LG, et al. Scene-Selective Cortical Regions in Human and Nonhuman Primates. J Neurosci. 2011 Sep 28;31(39):13771–85.

69. Awh E, Armstrong KM, Moore T. Visual and oculomotor selection: links, causes and implications for spatial attention. Trends in Cognitive Sciences. 2006 Mar 1;10(3):124–30.

70. Majka P, Bai S, Bakola S, Bednarek S, Chan JM, Jermakow N, et al. Open access resource for cellular-resolution analyses of corticocortical connectivity in the marmoset monkey. Nat Commun. 2020 Feb 28;11(1):1133.

71. Schaeffer DJ, Klassen LM, Hori Y, Tian X, Szczupak D, Yen CCC, et al. An open access resource for functional brain connectivity from fully awake marmosets. NeuroImage. 2022 May 15;252:119030.

72. Belin P, Fecteau S, Bédard C. Thinking the voice: neural correlates of voice perception. Trends in Cognitive Sciences. 2004 Mar 1;8(3):129–35.

73. Bodin C, Trapeau R, Nazarian B, Sein J, Degiovanni X, Baurberg J, et al. Functionally homologous representation of vocalizations in the auditory cortex of humans and macaques. Current Biology. 2021 Nov 8;31(21):4839–4844.e4.

74. Miller CT, Freiwald WA, Leopold DA, Mitchell JF, Silva AC, Wang X. Marmosets: A Neuroscientific Model of Human Social Behavior. Neuron. 2016 Apr 20;90(2):219–33.

75. Liu C, Yen CCC, Szczupak D, Ye FQ, Leopold DA, Silva AC. Anatomical and functional investigation of the marmoset default mode network. Nat Commun. 2019 Apr 29;10(1):1975.

76. Raichle ME, MacLeod AM, Snyder AZ, Powers WJ, Gusnard DA, Shulman GL. A default mode of brain function. Proceedings of the National Academy of Sciences. 2001 Jan 16;98(2):676–82.

77. Watanabe M. Are there internal thought processes in the monkey?—Default brain activity in humans and nonhuman primates. Behavioural Brain Research. 2011 Aug 1;221(1):295–303.

78. Mantini D, Gerits A, Nelissen K, Durand JB, Joly O, Simone L, et al. Default Mode of Brain Function in Monkeys. J Neurosci. 2011 Sep 7;31(36):12954–62.

79. Hutchison RM, Everling S. Monkey in the middle: why non-human primates are needed to bridge the gap in resting-state investigations. Front Neuroanat [Internet]. 2012 Jul 26 [cited 2024 Apr 15];6. Available from: https://www.frontiersin.org/articles/10.3389/fnana.2012.00029

80. Stafford JM, Jarrett BR, Miranda-Dominguez O, Mills BD, Cain N, Mihalas S, et al. Large-scale topology and the default mode network in the mouse connectome. Proceedings of the National Academy of Sciences. 2014 Dec 30;111(52):18745–50.

81. Lu H, Zou Q, Gu H, Raichle ME, Stein EA, Yang Y. Rat brains also have a default mode network. Proceedings of the National Academy of Sciences. 2012 Mar 6;109(10):3979–84.

82. Liska A, Galbusera A, Schwarz AJ, Gozzi A. Functional connectivity hubs of the mouse brain. NeuroImage. 2015 Jul 15;115:281–91.

83. Abell F, Happé F, Frith U. Do triangles play tricks? Attribution of mental states to animated shapes in normal and abnormal development. Cognitive Development. 2000 Jan 1;15(1):1–16.

84. Castelli F, Happé F, Frith U, Frith C. Movement and Mind: A Functional Imaging Study of Perception and Interpretation of Complex Intentional Movement Patterns. In: Social Neuroscience. Psychology Press; 2004.

85. Wong RK, Selvanayagam J, Johnston KD, Everling S. Delay-related activity in marmoset prefrontal cortex. Cereb Cortex. 2023 Mar 21;33(7):3523–37.

86. Riley MR, Constantinidis C. Role of Prefrontal Persistent Activity in Working Memory. Front Syst Neurosci [Internet]. 2016 Jan 5 [cited 2024 Jul 4];9. Available from: https://www.frontiersin.org/journals/systems-neuroscience/articles/10.3389/fnsys.2015.00181/full

87. Belcher AM, Yen CC, Stepp H, Gu H, Lu H, Yang Y, et al. Large-Scale Brain Networks in the Awake, Truly Resting Marmoset Monkey. J Neurosci. 2013 Oct 16;33(42):16796–804.

88. Ngo GN, Hori Y, Everling S, Menon RS. Joint-embeddings reveal functional differences in default-mode network architecture between marmosets and humans. NeuroImage. 2023 May 15;272:120035.

89. Ma L, Selvanayagam J, Ghahremani M, Hayrynen LK, Johnston KD, Everling S. Single-unit activity in marmoset posterior parietal cortex in a gap saccade task. Journal of Neurophysiology. 2020 Mar;123(3):896–911.

90. Ghahremani M, Johnston KD, Ma L, Hayrynen LK, Everling S. Electrical microstimulation evokes saccades in posterior parietal cortex of common marmosets. Journal of Neurophysiology. 2019 Oct;122(4):1765–76.

91. Hardee JE, Thompson JC, Puce A. The left amygdala knows fear: laterality in the amygdala response to fearful eyes. Social Cognitive and Affective Neuroscience. 2008 Mar 1;3(1):47–54.

92. Wright CI, Fischer H, Whalen PJ, McInerney SC, Shin LM, Rauch SL. Differential prefrontal cortex and amygdala habituation to repeatedly presented emotional stimuli. NeuroReport. 2001 Feb 12;12(2):379–83.

